# Single-cell RNA transcriptomics identifies Hivep3 as essential in regulating the development of innate-like T lymphocytes

**DOI:** 10.1101/2020.06.08.135129

**Authors:** Sai Harsha Krovi, Jingjing Zhang, Mary Jessamine Michaels-Foster, Tonya Brunetti, Liyen Loh, James Scott-Browne, Laurent Gapin

## Abstract

Most T lymphocytes leave the thymus as naïve cells with limited functionality. However, unique populations of T cells, commonly known as innate-like T cells, differentiate into functionally distinct effector subsets during thymic development under the influence of the transcription factor PLZF. Here, we profiled >10,000 differentiating thymic iNKT cells using single-cell RNA sequencing to provide a comprehensive transcriptional landscape of their maturation, function, and fate decision in steady state. We identified *Hivep3*, a zinc finger transcription factor and adaptor protein, as a key factor that is expressed in early precursors and regulates the post-selection proliferative burst, differentiation and functions of iNKT cells. Importantly, we extended these results to other PLZF^+^ innate-like T cell populations, highlighting the unique and common requirement of *Hivep3* to the development of all innate-like T cells.

## Introduction

Two fundamental features of the adaptive T cell response are its enormous antigen recognition capacity and its ability to form immunological memory. The former endows T cells with a potential repertoire capable of targeting the myriad pathogens a host might encounter while the latter helps the host respond with significantly faster kinetics in the event of a reinfection with a previously encountered pathogen. Although the vast majority of T cells fit this paradigm, recent work has begun to highlight the importance of innate-like T (T_inn_) cells, which differ both in their phenotype and response kinetics. T_inn_ cells, such as invariant natural killer T (iNKT) cells and mucosal-associated invariant T (MAIT) cells, tend to establish permanent residency in tissues, express restricted T cell antigen receptor (TCR) repertoires and share a non-MHC class I or II restriction requirement for antigen recognition ^
1,2^. Compared to their conventional T cell (T_conv_) counterparts, T_inn_ cells are also not constrained by the need for prior sensitization and exist in a pre-primed “memory” state, ready to respond within minutes of stimulation, despite being antigen-inexperienced ^3^. This is due to the fact that unlike T_conv_ cells, T_inn_ cells acquire an effector and memory phenotype over the course of their development in the thymus ^4^.

Thymic development of iNKT cells has been extensively studied over the past decade. Although they share their developmental origins with T_conv_ up to the double positive (DP) thymocyte stage, their paths then diverge ^5^. While DP precursors for T_conv_ are selected on peptide-MHC complexes expressed on thymic epithelial cells, iNKT DP precursors are selected on lipids bound to the MHC-I-like molecule (MHC-Ib) CD1d expressed on neighboring DP thymocytes ^6^. These DP-DP synapses promote commitment to the iNKT lineage by inducing high levels of the transcription factor Egr2, which in turn lead to expression of the lineage-determining BTB-ZF transcription factor PLZF ^7^. PLZF subsequently directs the further differentiation of iNKT cells into three distinct mature effector subsets, iNKT1, iNKT2 and iNKT17, analogous to the CD4^+^ T cell polarized subsets observed in the periphery. Also similar to the CD4^+^ T cell subsets, each of these iNKT subsets expresses the master transcription factors that drive their fates, with iNKT1 cells expressing T-bet, iNKT17 expressing Rorγt, and iNKT2 expressing high levels of PLZF and GATA3 (Ref ^4,8^). Thus, iNKT effector subsets arise as a consequence of their thymic development at steady-state instead of differentiation in peripheral tissues due to inflammatory cues. Additionally, some iNKT subsets have been shown to secrete cytokines at steady-state, thereby influencing their surrounding tissue microenvironments in non-redundant manners ^8^. More recently, MAIT cells were shown to follow a similar developmental pathway ^9,10^, with the key distinguishing feature being that they are selected by other DP cells expressing the MHC-Ib molecule MR1 presenting vitamin metabolites ^11^.

While a large body of knowledge is available on iNKT cell characteristics and development in the thymus and in the periphery ^12^, the developmental steps underlying iNKT cell differentiation, and by extension MAIT cells, remain, nevertheless, incomplete. In particular, studies have largely focused on analyzing iNKT cells in bulk, potentially missing developmental intermediate populations. To overcome this limitation, we characterized the transcriptomes of thymic iNKT cells at the single cell level (scRNA-seq) to reveal the underlying heterogeneity that exists over the course of iNKT cell development. We performed scRNA-seq on a large number of purified thymic iNKT cells (>10,000 cells) and established a high-resolution outline of iNKT cell development in an unbiased fashion. Our results illustrated the transcriptional signature of the previously identified stage 0, iNKT1, iNKT2 and iNKT17 cells, and further exposed heterogeneity amongst iNKT1 cells. In addition, we identified the *Hivep3 gene*, a member of the human immunodeficiency virus type enhancer-binding protein family, which encodes a large protein that can function both as an adaptor ^13,14^ and can bind NF-κB motifs in the genome to control transcriptional programs ^15,16^. *Hivep3* is highly and transiently expressed in early stage 0 iNKT cell precursors as well as in the earliest MAIT cell progenitors. Owing to an early block in iNKT cell development at the CD24^high^ stage, *Hivep3*-deficient mice exhibited a profound loss of both iNKT and MAIT cells in all examined tissues, with the few remaining cells being functionally impaired. The frequency and number of PLZF^+^Vγ1^+^Vδ6.3^+^ γδ T cells, another T_inn_ cell population ^17^, were also largely reduced in KO mice. Furthermore, scRNA-seq of *Hivep3*^−/−^ thymic iNKT cells revealed a crucial role for *Hivep3* in regulating the expression of several key developmental genes, including *Hdac7*, *Drosha* and *Zbtb16* (the gene coding for PLZF). ATAC-seq analysis of mature iNKT cell subsets showed that in absence of *Hivep3*, chromatin accessibility was increased at regions enriched for NF-κB binding motifs. Altogether, our results identify *Hivep3* as a key regulator that controls, transcriptionally and post-transcriptionally, the development of naïve versus PLZF^+^ innate T lymphocytes.

## Results

### Single-cell RNA-seq of thymic C57BL/6 iNKT cells

To investigate the transcriptional landscape of iNKT cell differentiation in the mouse thymus, we performed droplet-based single cell RNA-seq (scRNA-seq) analysis of PBS57-CD1d-tetramer^+^ cells sorted from the thymi of 8-week old female C57BL/6 mice using the Chromium system (10x Genomics). To enrich for early precursors and developmental intermediates, CD1d-tetramer^+^ cells were sorted at a 50:50 ratio of CD44^low^ to CD44^high^ cells. Two independent experiments were performed. The two distinct sc-RNAseq datasets were next integrated and corrected for batch effects into an integrated reference dataset using the fastMNN algorithm ^18^. After quality control steps aimed at removing low-quality cells with few genes and/or stressed/apoptotic cells with high expression of mitochondrial genes and a filtering step to remove multiplets, a total of 10,506 cells (6937 cells from the first sample and 3569 cells from the second sample) were retained for further analysis. We used uniform manifold approximation and projection (UMAP) for dimensionality reduction to display all iNKT cells in a shared low-dimensional representation (**Fig 1a**). To further characterize the subpopulation structures, we applied unsupervised graph-based clustering, which yielded 11 distinct clusters (**Fig 1b**), with an average of 1956 genes and 6617 UMI expressed by the cells in each cluster (**Fig 1c**). The expression of *Cd24a*, *Cd69*, *Egr2*, *Il2rb*, *Cd44*, *Zbtb16* (encoding PLZF), *Rorc* (encoding Rorγt), *Tbx21* (encoding T-bet), *Gata3* (encoding GATA3), *Ifng*, *Il4*, *Ccr7*, *Ccr6*, *Il17rb* and *Cd4* were used to define immature populations and effector subsets (**Fig 1d**). iNKT17 cells (*Rorc*^+^, *Ccr6*^+^, *Zbtb16*^+^, *Cd4*^−^) were found in a single cluster (cluster 5), while iNKT1 cells (*Il2rb*^+^, *Tbx21*^+^, *Ifng*^+^) were found in five clusters (clusters 6 to 10). iNKT2 cells (*Zbtb16*^+^, *Gata3*^+^, *Cd4*^+^, *Il4*^+^) were contained within 4 clusters (clusters 1 to 4), with three of these clusters (clusters 1, 2 and 3) corresponding to cycling cells (**Fig 1f and Fig 2c, d**). The earliest iNKT precursor cells, stage 0 cells, were assigned to cluster 0. Expression of *Cd24a*, the proliferation marker *Ki67*, and the master transcription factors defining each of the mature iNKT cell subsets (*Zbtb16*, *Rorc* and *Tbx21*) confirm these identities (**Fig 1f**), with the top five genes that characterize each of these clusters shown in **Fig 1e**.

**Figure 1.**
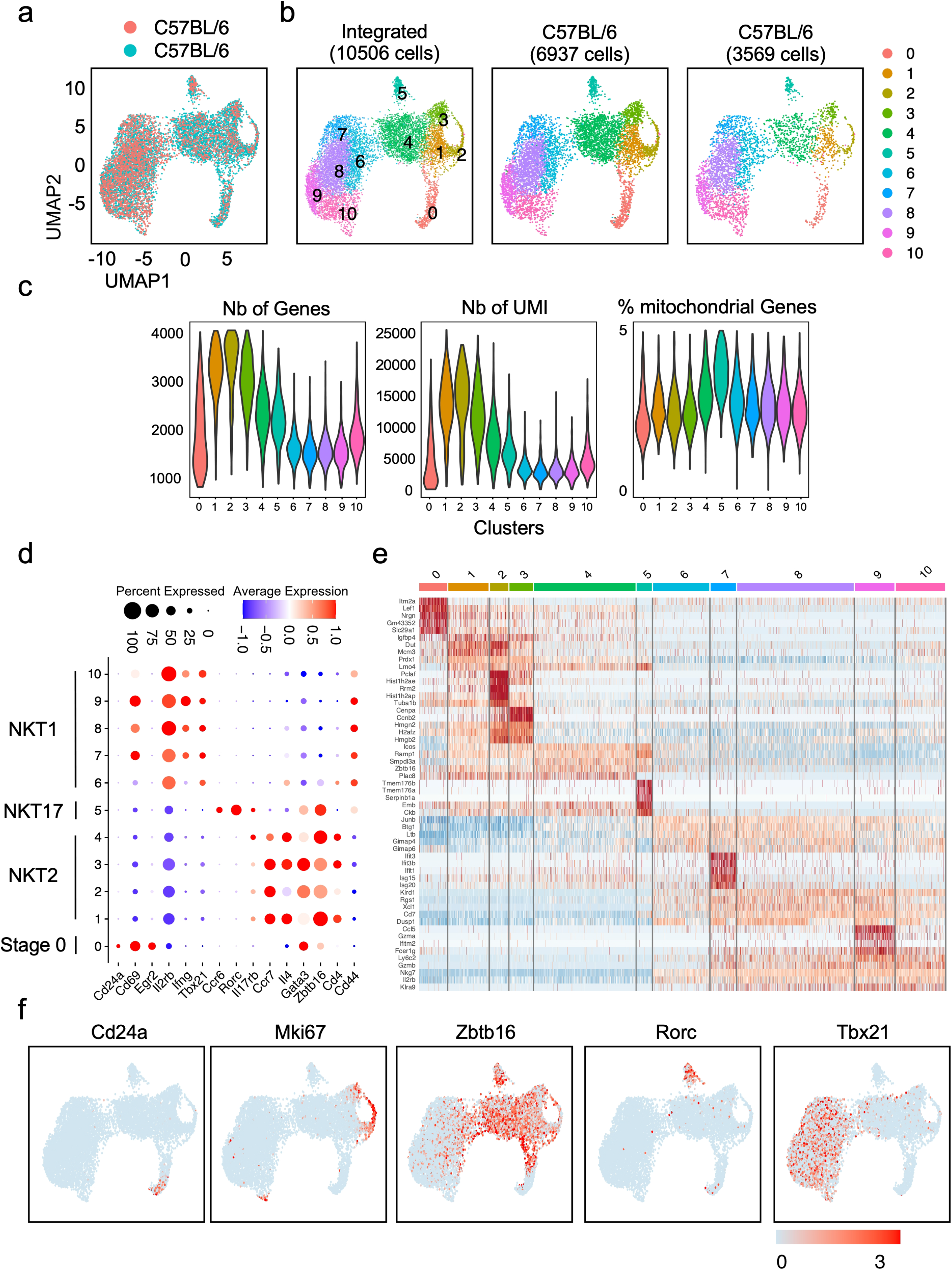
ScRNA-seq analysis of steady-state C57BL/6 thymic iNKT cells. (**a**) Uniform manifold approximation and projection (UMAP) of two independent scRNA-seq data sets from C57BL/6 thymic iNKT cells integrated using FastMNN and colored by sample of origin. (**b**) UMAP of 10,506 iNKT cells colored by inferred cluster identity. (**c**) Violin plot depicting the number of gene, number of UMI and percentage of mitochondrial genes expressed in each cluster (**d**) Dot plot showing scaled expression of selected signature genes for each cluster colored by average expression of each gene in each cluster scaled across all clusters. Dot size represents the percentage of cells in each cluster with more than one read of the corresponding gene. (**e**) Heatmap showing row-scaled expression of the 5 highest differentially expressed genes (DEGs, Bonferroni-corrected P-values < 0.05, Wilcoxson-test) per cluster for all iNKT cells. (**f**) Expression of five typical genes used to define most common iNKT cell subsets and cycling cells.

**Figure 2.**
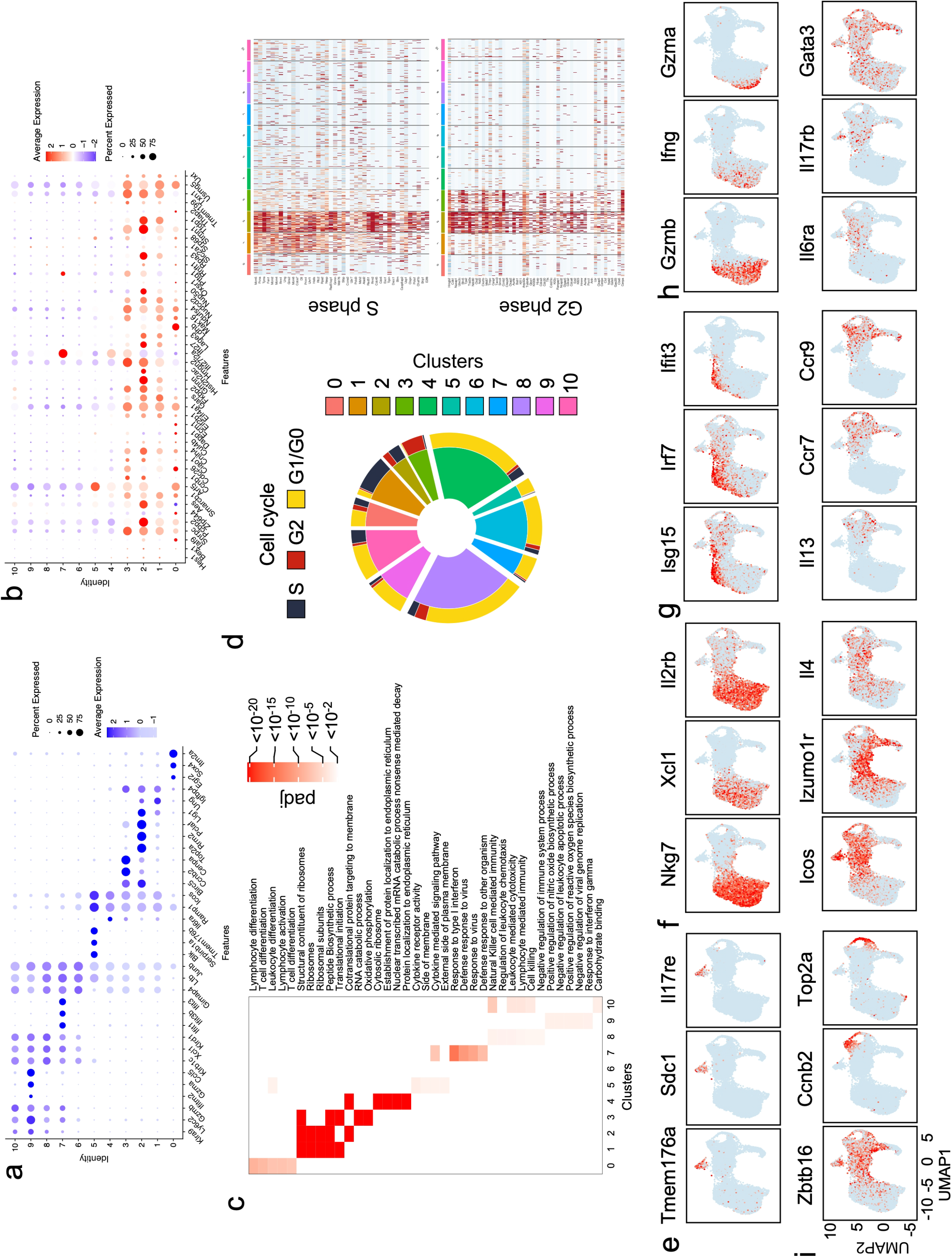
Thymic iNKT subsets display distinct gene expression profiles. (**a**) Dot plot showing scaled expression of the top 3 highest DEGs for each cluster colored by average expression of each gene in each cluster scaled across all clusters. Dot size represents the percentage of cells in each cluster with more than one read of the corresponding gene. (**b**) Dot plot showing scaled expression of the 44 genes previously described as the iNKTp signature, for each cluster colored by average expression of each gene in each cluster scaled across all clusters. Dot size represents the percentage of cells in each cluster with more than one read of the corresponding gene. (**c**) Gene ontology (GO) analysis of DEGs for each cluster. Selected GO terms with Benjamini-Hochberg-corrected P-values < 0.05 (one-sided Fisher's exact test) are shown. Heatmaps display p_adj_ value significance of enrichment of GO terms in each cluster. (**d**) Pie plot depicting the proportion of cells in each cluster and for each cluster the proportion of cells in each phase of the cell cycle. Heatmap showing row-scaled expression of cell cycle-related genes for each cluster. (**e**) Expression of three typical genes used to define iNKT17 cells. (**f**) Expression of three typical genes used to define iNKT1 cells. (**g**) Expression of three typical genes used to define interferon signaling genes. (**h**) Expression of three typical genes used to define cells in cluster 9. (**i**) Expression of twelve genes used to illustrate and define the diversity amongst iNKT2/cycling cells.

### Transcriptional heterogeneity of thymic iNKT cells

We next identified the cluster-specific signature genes (**Fig 2a** **and Sup. Table I**), performed gene ontology analysis associated with gene expression in each cluster (**Fig 2c**) and determined the cell cycle status of the cells (**Fig 2d**). Due to our sorting strategy that enriched for immature CD44^low^ cells, we identified 565 stage 0 cells (cluster 0), allowing for a more in-depth analysis of their transcriptional profile than previously achieved ^19^. Differentially expressed genes (DEG) in cluster 0 comprised several genes associated with lymphocyte activation and T cell differentiation (**Fig 2c**), including *Itm2a* ^20^, *Pdcd1* (encoding PD-1), *Cd27*, *Cd28*, *Cd69*, *Slamf6, Cd81, Ldhb* and *Bcl2* (coding for the anti-apoptotic protein Bcl-2) (**Fig 3a**). In agreement with previous flow cytometry results ^21^, stage 0 iNKT cells did not express detectable *Cd4* or *Cd8a* transcripts (**Fig 3a**). Several genes coding for transcription factors, such as *Lef1* (ref ^22^), *Sox4* (ref ^23^), *Egr2* and *Egr1* (ref ^7^), *Tox* ^24^, *Id3* (ref ^25^), *Myb* ^26^, and *Ikzf1* (coding for Helios), including some that have been previously implicated in iNKT cell development, were highly expressed in stage 0 iNKT cells (**Fig 3a**). High expression of the protein products encoded by some of these genes in stage 0 iNKT cells was verified by flow cytometry, thereby validating the scRNA-seq data (**Fig 3c**). Interestingly, we also detected high levels of *Hivep3* transcripts (**Fig 3a**), a gene previously known to be induced upon CD3/CD28 activation of CD4 T cells ^14^, and proposed to be a potential Egr2 target in the early stages of development of iNKT cells 7.

**Figure 3.**
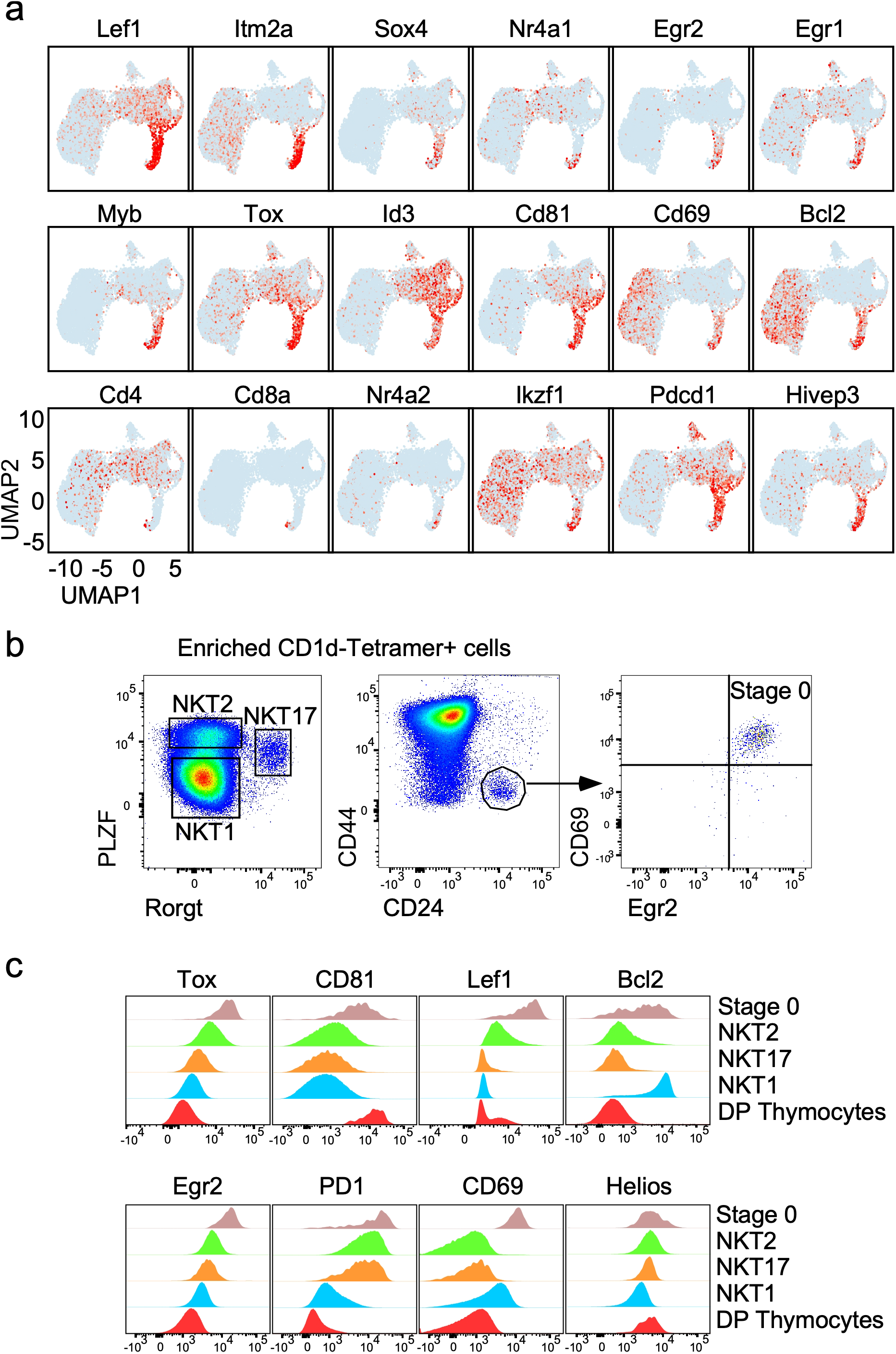
Gene expression profile of stage 0 iNKT cells. (**a**) Expression patterns of eighteen stage 0 iNKT-specific genes in our scRNA-seq data. Each dot represents one cell and gene expression is plotted along a colorimetric gradient, with red corresponding to high expression. (b) Gating strategy to identify stage 0, iNKT1, iNKT2 and iNKT17 cell subsets by flow cytometry. (c) Histograms displaying protein expression levels by flow cytometry within each iNKT subset for the indicated proteins. Protein expression in DP thymocytes is also displayed as a control.

A multi-potent CCR7^+^ iNKT cell progenitor (iNKTp), with a unique transcriptional signature has been described as a developmental intermediary following commitment to the iNKT cell lineage by stage 0 cells ^27^. However, we could not assign this gene signature, composed of 44 different genes, to a transcriptionally unique set of cells and instead observed this gene set expressed by cells in multiple clusters (Clusters 1, 2 and 3) (**Fig 2b**). Cells within these clusters have upregulated expression of *Zbtb16* (**Fig 1d, e** **and** **f**) as well as several other markers usually associated with the iNKT2 phenotype, including *Icos* and *Izumo1r* (**Fig 2i**). These cells have high ribosomal and mitochondrial activity (**Fig 2c**), in agreement with their high proliferative status (**Fig 2d, i**). Computation of cell cycle scores revealed that cells in cluster 1 were predominantly in S phase, cells in cluster 2 comprised S and G2 phases, and cells in cluster 3 largely belonged to the G2 phase of the cell cycle (**Fig 2d**). These results are in agreement with the proliferative burst known to occur after stage 0 in iNKT cells ^28^. Cells in cluster 4 had a similar iNKT2 transcriptional profile (**Fig 2i**), but with increased expression of *Il6ra*, *Il17rb*, *Icos*, *Plac8* and *Il4* transcripts, decreased expression of transcripts encoding for *Ccr7*, *Ccr9*, *Il13* and undetectable expression of cell-cycle genes (**Fig 2i, d** **and Sup. Table I**).

Cells pertaining to cluster 5 expressed high levels of the master regulator *Rorc* (**Fig 1f**) and other genes whose expression have been shown to be directly dependent on Rorgt in T_H_17 cells ^29^, including *Tmem176a*/*b*, *Ltb4r1*, *Il1r1*, and *Il23r* (**Fig 2e** **and Sup. Table I**). iNKT17 signature genes also comprised several transcripts whose products are involved in tissue residency (*Itgb7*, *Ccr6*, *Aqp3*) as well as several additional cytokine and cell signaling receptors (*Il17re*, *Il18r1*, *Il7r*, *Sdc1*) (**Fig 2e and Sup. Table I)**and were defined by their cytokine receptor and signaling pathway activity (**Fig 2c**).

Cells in the five remaining clusters (clusters 6, 7, 8, 9 and 10) could all be considered iNKT1 cells, as they expressed transcripts for *Tbx21* (**Fig 1f**), *Il2rb* (coding for CD122), *Nkg7*, and *Xcl1* (**Fig 2f**). Interestingly, cells in cluster 6, which neighbor the iNKT2 cells belonging to cluster 4 on the UMAP (**Fig 1**), appeared as a transitional population bearing a mixed transcriptional signature observed in cells from cluster 8 (*Klrd1*, *Xcl1*, *Cd7*, *Nkg7*, *Ms4a4b*, *Dusp1* and *Dusp2*) and cells from cluster 4 (*Izumo1r*) (**Fig 2f**). These cells were nevertheless further defined by higher expression of transcripts encoding for *Junb*, *Ltb* and several members of the GTPase of the immunity-associated protein (GIMAP) family (*Gimap4, 3 and 6*) (**Sup. Table I)**. These results allude to the possibility that cells in cluster 6 could represent an intermediary state between iNKT2 cells and iNKT1 cells. Independent pseudo-time ordering using the Monocle v3 (ref ^30^) and Slingshot ^31^ trajectory inference packages to place iNKT cell populations along possible developmental trajectories, with stage 0 cells defined as the root, supports this possibility (**Sup. Fig 1**). Cells in clusters 9 and 10 had enriched expression of transcripts for *Gzma*, *Gzmb*, *Ifng*, *Ccl5*, *Xcl1*, *Fcer1g* and *Ly6c2* and killer cell lectin type receptors (*Klrk1*, *Klre1*, *Klra5*, *Klra9*, *Klrc1*, *Klrd1*, *Klrc2*) suggesting that they might represent a further step in the maturation of iNKT1 cells associated with the acquisition of cytotoxicity (**Fig 2e** **and Sup. Table I**). Finally, cells in cluster 7 had a unique transcriptional signature with high expression of type I interferon response genes such as *Isg15* and *20*, *Ifit1* and *3*, *Irf7* and *Stat1* (**Fig 2c, g** **and Sup. Table I)**, perhaps suggesting that tonic interferon signaling might be involved in the final maturation of iNKT cells, similar to what has been reported for conventional T cell development^32^.

### Altered iNKT cell development in Hivep3 deficient mice

To investigate the function of *Hivep3* in iNKT cells, we analyzed Hivep3^−/−^ mice backcrossed to the C57BL/6 background ^33^. *Hivep3*^−/−^ mice exhibited a large reduction in the proportion and numbers of iNKT cells in all organs examined, including the thymus, spleen, liver, inguinal lymph nodes and lungs (**Fig 4a** **and** **b**). This lack of iNKT cells was a cell-intrinsic consequence of Hivep3 deficiency as revealed by competitive bone marrow chimeras (**Fig 4c**), in which only the wildtype bone marrow cells were able to efficiently reconstitute the thymic iNKT cell compartment while conventional CD4^+^ single positive thymocytes were equivalently reconstituted by both wildtype and *Hivep3*^−/−^ bone marrow cells (**Fig 4c**), as expected by the normal development of conventional T cells observed in *Hivep3*^−/−^ mice (**Fig 4h**). To dissect the developmental stage at which the absence of *Hivep3* expression affected iNKT cells, we performed magnetic bead-based tetramer enrichment of thymic iNKT cells and stained for markers used to define the various stages of development (CD24, CD44 and NK1.1) (**Fig 4d**) as well as for transcription factors used to define iNKT cell subsets (PLZF, Rorγt and T-bet) (**Fig 4f**). Our analysis revealed that the percentages of cells at stages 0, 1 and 2 were increased in the absence of *Hivep3* while the proportion of cells at stage 3 was decreased. However, while the total cell number of stage 0 iNKT cells was not significantly different between C57BL/6 and *Hivep3*^−/−^ mice, the number of cells at any subsequent developmental stages was largely and significantly decreased (**Fig 4e**). These results indicate that positive selection of iNKT cells is not altered in the absence of Hivep3 expression, but that a developmental block occurs immediately following stage 0 (**Fig 3a**). Surprisingly, the relative proportions of iNKT2, iNKT17 and iNKT1 were only minimally affected, although the numbers of all mature iNKT subsets were dramatically reduced in *Hivep3*^−/−^ thymi (**Fig 4f** **and** **g**). Together, these results suggest that although the development of iNKT cells in *Hivep3*^−/−^ mice is essentially arrested at stage 0, a few cells can overcome this block and differentiate into the mature iNKT cell subsets. Nevertheless, we noticed that the remaining iNKT cells in *Hivep3*^−/−^ mice had overall higher TCR expression levels (**Fig 4a, d**) and that PLZF protein levels in each of the iNKT subsets were reduced (**Fig 4f**), collectively hinting at the possibility that the few iNKT remnants in *Hivep3*^−/−^ mice did not develop normally.

**Figure 4.**
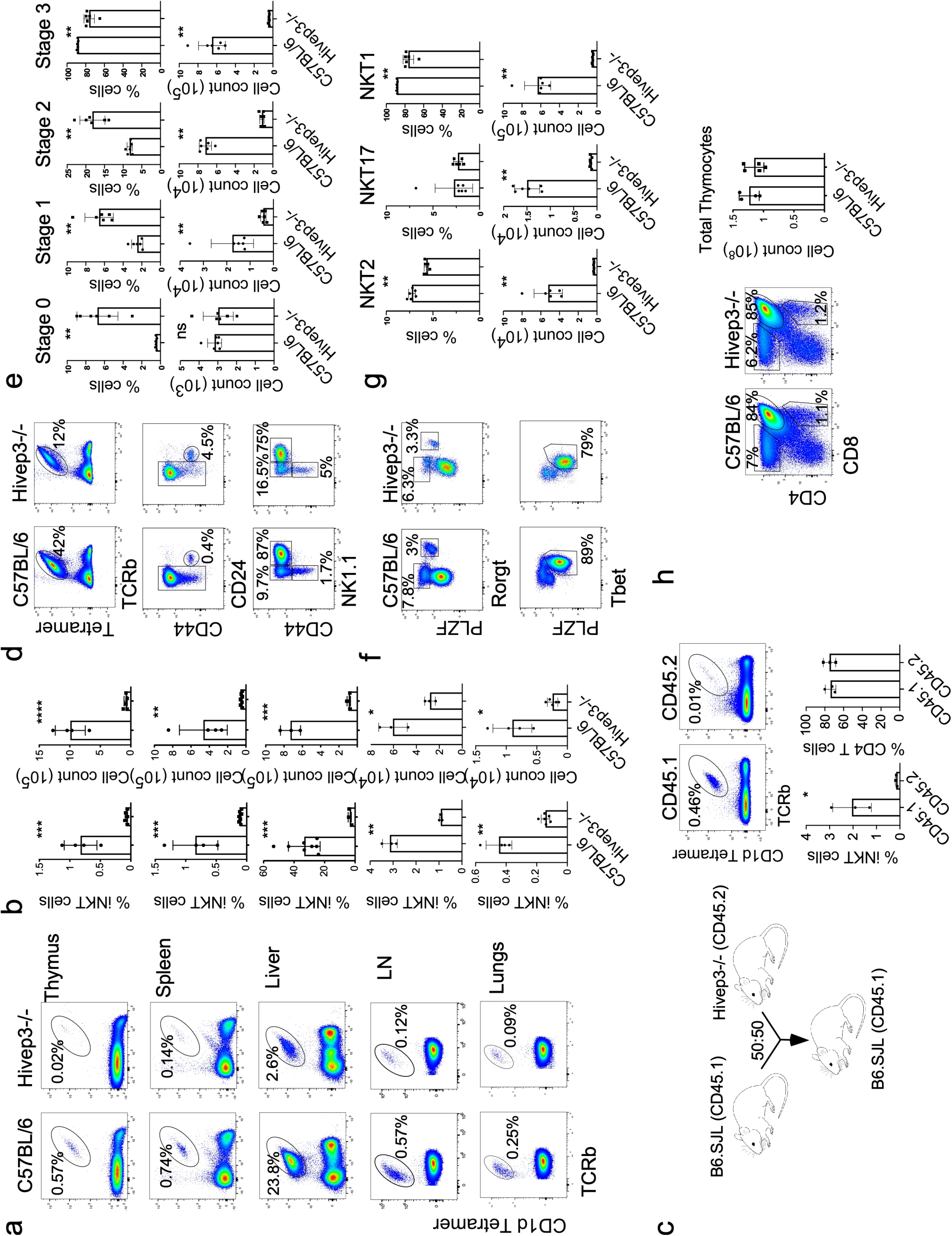
Intrinsic deficiency of iNKT cells in *Hivep3*^−/−^ mice. (**a**) iNKT cell proportions in C57BL/6 and *Hivep3*^−/−^ mice were identified in the indicated tissues by staining with PBS57-CD1d tetramers and an antibody targeting TCRβ. (**b**) Quantified iNKT cell proportions and numbers in C57BL/6 and *Hivep3*^−/−^ mice in each of the indicated tissues. (**c**) *Left*, Cartoon depicting the competitive bone marrow chimera strategy in which CD45.1 C57BL/6 bone marrow was mixed at a 50:50 ratio with CD45.2 *Hivep3*^−/−^ bone marrow and subsequently transferred into irradiated CD45.1 recipient mice. *Right*, iNKT and single positive CD4^+^ T cell proportions were quantified from both C57BL/6 (CD45.1) and *Hivep3*^−/−^ (CD45.2) mice after immune reconstitution. (**d, e**) iNKT cells from C57BL/6 and *Hivep3*^−/−^ mice were categorized based on expression of CD24, CD44, NK1.1 into stage 0, stage 1, stage 2 and stage 3 and subsequently quantified for their relative proportions and numbers. (**f, g**) iNKT cells from C57BL/6 and *Hivep3*^−/−^ mice were categorized based on expression of PLZF, Rorγt and T-bet into iNKT1, iNKT2 and iNKT17 and subsequently quantified for their relative proportions and numbers. (**h**) *Left*, Flow cytometry analysis depicting DP, CD4 and CD8 proportions in WT and KO mice. *Right*, Histograms quantifying total thymic cellularity.

### ScRNA-seq analysis of Hivep3^−/−^ iNKT cells

To gain further insight into how *Hivep3* deficiency affects iNKT cell development, we performed two independent scRNA-seq experiments using PBS57-CD1d-tetramer^+^ cells sorted from the thymi of 8-week old *Hivep3*^−/−^ mice, using a similar sort strategy to the one we utilized to sort our C57BL/6 samples (**Fig 1**). The two distinct scRNA-seq datasets were integrated with the C57BL/6 data and corrected for batch effects (**Fig 5a, b**). The transcriptional profile of *Hivep3*^−/−^ iNKT cells was rather similar to C57BL/6 iNKT cells, as we had previously observed using flow cytometry **(Fig 4)**, with unsupervised graph-based clustering revealing 10 different clusters common to both genotypes (**Fig 5b**). The proportion of cells in cluster 0 (stage 0 cells) was increased in Hivep3^−/−^ iNKT cells, in agreement with a block in development at, or immediately following, that stage (**Fig 4**). A greater proportion of cells from *Hivep3*^−/−^ than from C57BL/6 mice was also found in cluster 1 (~1% of C57BL/6 cells vs ~7.5% of *Hivep3*^−/−^ cells). Interestingly, these *Hivep3*^−/−^ cells displayed a predominant iNKT1 gene signature (**Sup Fig 2**). By contrast, the proportion of proliferating cells in clusters 2 and 3 was under-represented in *Hivep3*^−/−^ cells compared to C57BL/6 cells (**Fig 5c**), which was confirmed by staining for the Ki67 proliferation marker and BrdU incorporation (**Sup Fig 3**). Although the proportion of cells in the iNKT2 and iNKT17 clusters (clusters 4 and 5) did not appear dramatically different between C57BL/6 and *Hivep3*^−/−^ cells, *Hivep3*^−/−^ iNKT1 cells were largely in cluster 9 and fewer cells were assigned to cluster 6 compared to the C57BL/6 iNKT1 cells (**Fig 5b** **and** **c**). Altogether, these results suggest that in the absence of *Hivep3*, iNKT development is blocked at stage 0, with a few cells able to further progress in an aberrant fashion, bypassing the proliferative burst and instead acquiring an iNKT1 transcriptional signature earlier during development.

**Figure 5.**
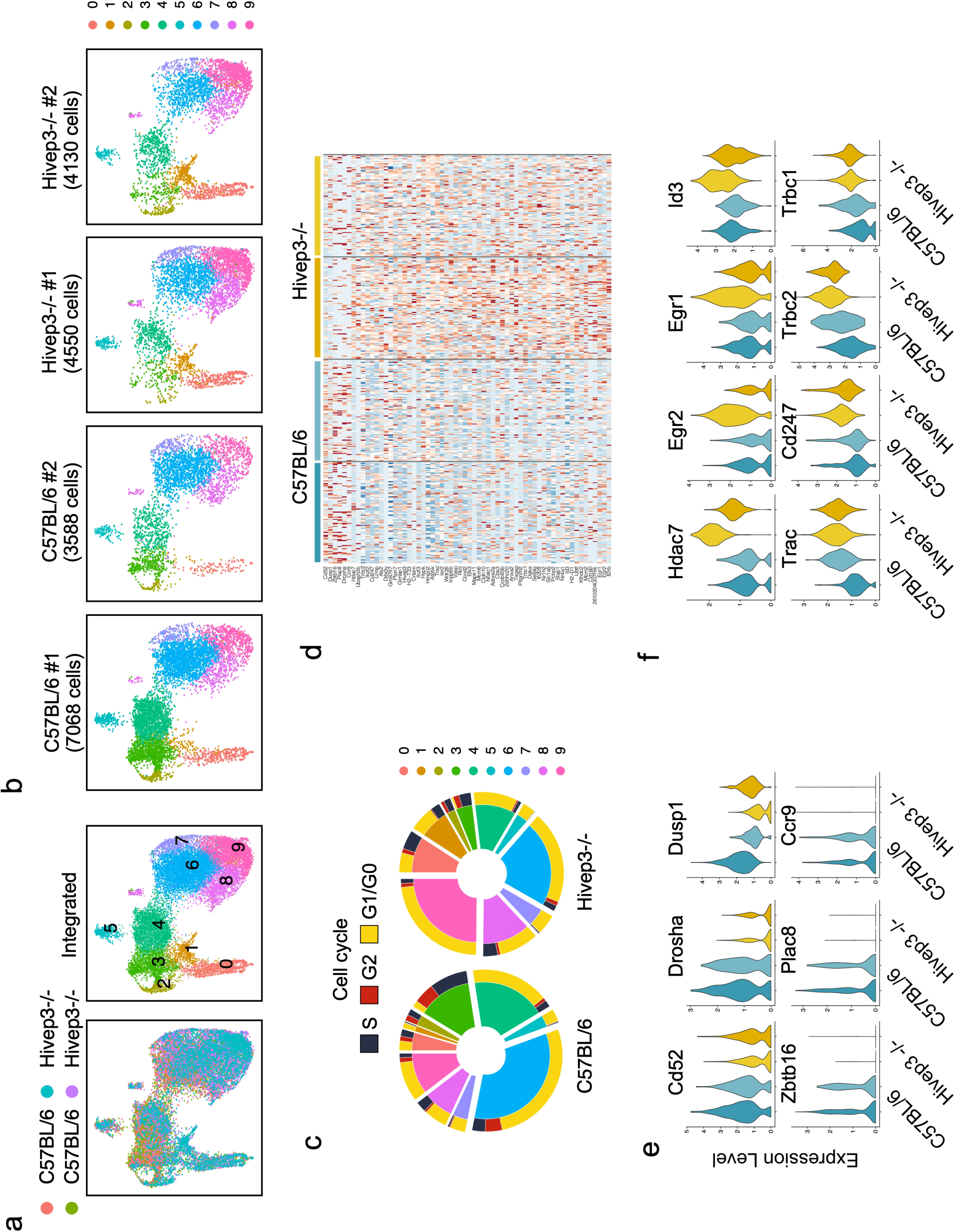
ScRNA-seq analysis of steady-state thymic iNKT cells from *Hivep3* ^−/−^ mice. (**a**) *Left*, Uniform manifold approximation and projection (UMAP) displaying two independent scRNA-seq data sets from C57BL/6 and two independent *Hivep3*^−/−^ thymic iNKT cells colored by sample of origin after MNN batch-correction. *Right*, UMAP colored by inferred cluster identity for the integrated dataset. (**b**) Individual UMAP plots for each biological sample in the dataset, with the sample and number of cells passing quality control and filtering indicated above each plot. (c) Cell cycle scores were computed for each cell in the dataset using the R Seurat package and then categorized into S, G2 or G1/G0 phases. Pie charts depict the proportion of C57BL/6 (*Left*) and *Hivep3*^−/−^ (*Right*) cells within each cluster belonging to a given phase of the cell cycle. (d) Heatmap displaying expression levels of 65 differentially expressed genes across stage 0 iNKT cells in the dataset. The sample to which a given cell belongs are indicated at the top of the heatmap. (**e, f**) Violin plots displaying aggregate expression levels per sample of genes that are expressed higher in C57BL/6 (**e**) or higher in *Hivep3*^−/−^ (**f**) stage 0 iNKT cells.

Since *Hivep3* expression in C57BL/6 cells is limited to the early stages of development (i.e. stage 0 cells, **Fig 3**), we next determined how its absence affected gene expression in stage 0 cells (**Sup. Table II**). We found 65 DEGs between C57BL/6 and *Hivep3*^−/−^ stage 0 cells (**Fig 5d**). Eight transcripts were expressed at lower levels in stage 0 Hivep3^−/−^ cells compared to C57BL/6 cells - *Cd52*, *Dusp1*, *Zbtb16*, *Plac8*, *Id2*, *Ccr9* and *Drosha* (**Fig 5d, e**). All other stage 0 DEGs had higher expression in Hivep3^−/−^ cells compared to C57BL/6 cells, including transcripts encoding the TCR chains and the CD3ζ subunit (*Trbc2*, *Trbc1*, *Trac*, *Cd247*) (**Fig 5f**), potentially explaining the higher level of TCR expression observed on *Hivep3*^−/−^ cells. Transcripts for several TFs downstream of TCR signaling, including *Egr1*, *Egr2*, *Nr4a1*, *Id3*, *Ikzf3* (**Fig 5f**), also displayed higher expression in *Hivep3*^−/−^ cells, suggesting that the KO cells might have received stronger signaling during positive selection. Interestingly, Hivep3^−/−^ stage 0 iNKT cells also expressed higher levels of transcripts encoding histone deacetylase 7 (*Hdac7*, **Fig 5f**), which was recently shown in a gain of function mouse transgenic model to similarly interfere with iNKT cell development ^34^.

Analyzing the remaining 9 clusters across strains revealed 785 DEGs between C57BL/6 and *Hivep3*^−/−^ cells (**Sup. Table II**). Notable amongst these DEGs were the decreased levels of expression in Hivep3^−/−^ cells compared to C57BL/6 cells of *Ccr9* and *Ccr7* transcripts, whose protein products are required for the migration of iNKT cell precursors from the cortex to the medulla ^35^. Transcripts encoding two Bcl-2 family members, *Bcl2a1b* and *Bcl2a1d*, were also largely decreased in Hivep3^−/−^ cycling cells, perhaps affecting their survival (**Fig 6** **and Sup. Table II**). *Cd4* and *Icos* transcripts were also decreased (**Fig 6**), which was also reflected at the protein level (**Sup. Fig 4**). Both *Il13* transcripts, which are found in cycling iNKT2 C57BL/6 cells and *Il4* transcripts, which are otherwise expressed in cycling and more mature iNKT2 cells, were reduced in the absence of *Hivep3* (**Fig 6**). These results suggest that *Hivep3*^−/−^ iNKT cells do not express appropriate amounts of the pre-formed mRNAs encoding these two cytokines, a hallmark of iNKT cell differentiation ^36,37^. Perhaps as a consequence, stimulation of peripheral iNKT cells *in vivo* with the agonist ligand α-GalCer, showed a defect in IL-4 production, while IFNγ production was less affected (**Sup Fig 5**). Similar to what was observed in progenitor cells (**Fig 5e**), the levels of *Zbtb16* transcripts were also found significantly decreased in subsequent developmental stages lacking *Hivep3* expression (**Fig 6**). Finally, expression of transcripts encoding the ribonuclease Drosha, which is responsible for the initiation step of miRNA processing, was mostly absent in Hivep3^−/−^ cells (**Fig 6**), perhaps contributing to the observation that 70% of DEGs in these 9 clusters display higher expression levels in *Hivep3*^−/−^ cells compared to C57BL/6 cells. These included several NKT1-associated transcripts, such as *Klra1* and *Klra5* (**Fig 6** **and Sup. Table II**).

**Figure 6.**
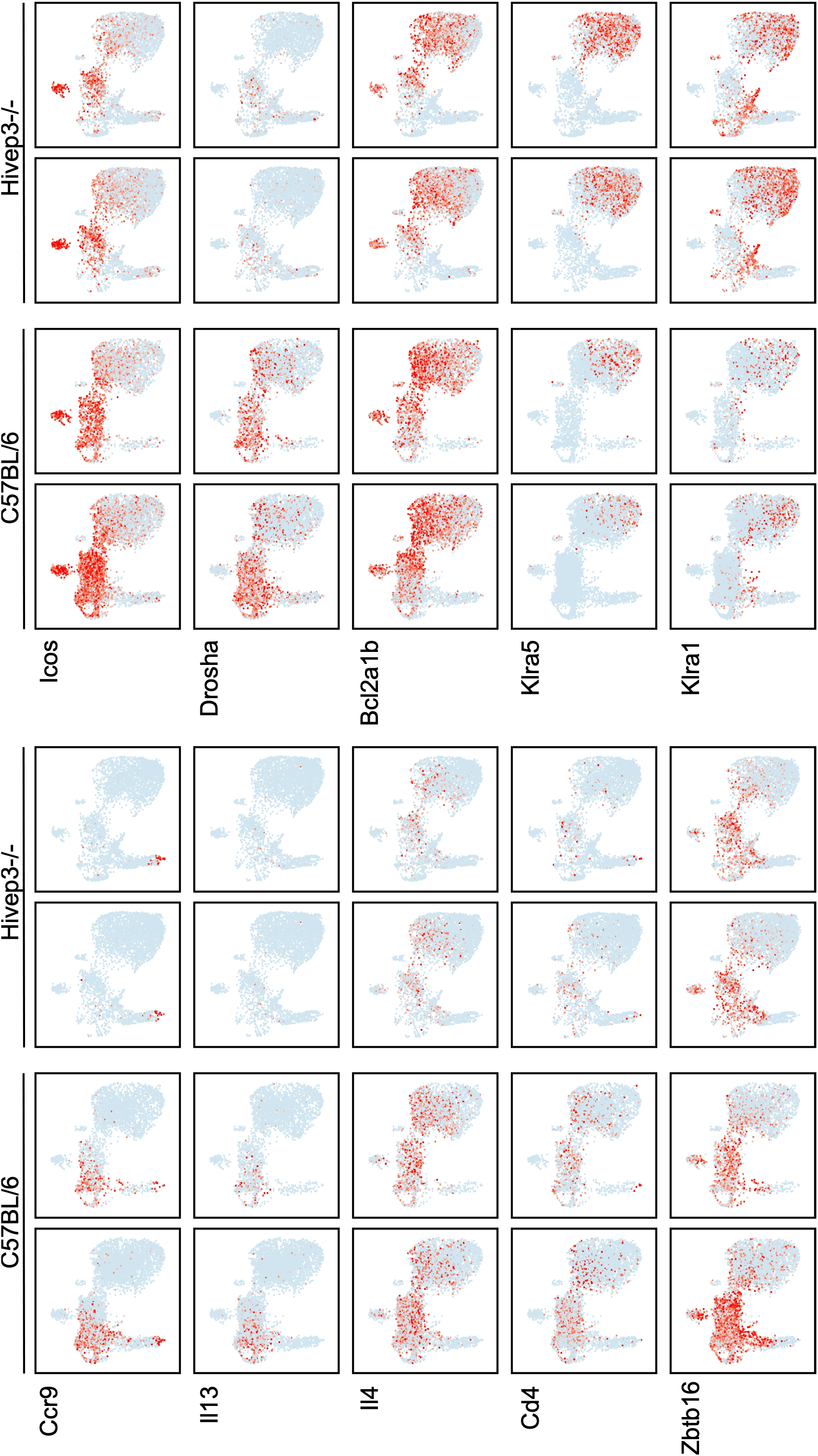
Expression profiles of genes differentially expressed between C57BL/6 and *Hivep3*^−/−^ iNKT cells by scRNA-seq. Expression of selected genes in the C57BL/6 and *Hivep3*^−/−^ single cell RNA-seq samples. Each dot represents one cell and gene expression is plotted along a colorimetric gradient, with red corresponding to high expression.

### Hivep3 activity influences chromatin landscape of iNKT subsets

To assess whether and how *Hivep3* might influence iNKT cell development via regulatory elements, we compared chromatin accessibility between C57BL/6 and *Hivep3*^−/−^ iNKT1, iNKT2 and iNKT17 cells with ATAC sequencing (ATAC-seq). iNKT1, iNKT2 and iNKT17 cell subsets each showed a unique chromatin landscape, with regions of accessibility specific to each subset (**Sup. Fig 6**), as previously shown ^21^. The analyses showed that *Hivep3*^−/−^ samples were closely related to the C57BL/6 sample for each iNKT cell subset, based on global chromatin acessibility signal compared by pairwise euclidean distances between each individual replicate (**Fig 7a**). To investigate whether there were specific changes in chromatin accessibility in iNKT cells from *Hivep3*^−/−^ mice, we compared the accessible regions between C57BL/6 and *Hivep3*^−/−^ backgrounds for each subset (**Fig 7b**). We identified 498 regions in iNKT1 cells and 730 regions in iNKT2 that were significantly (3 fold change, p_adj_<0.1) differentially accessible between C57BL/6 and *Hivep3*^−/−^ cells. In contrast, only 102 regions were found differentially accessible between C57BL/6 and *Hivep3*^−/−^ iNKT17 cells. Some, but not all, of these differentially accessible regions surrounded genes that were also found diffentially expressed transcriptionally between C57BL/6 and *Hivep3*^−/−^ cells. For example, the chromatin surrounding the *Cd247* gene was clearly more accessible in *Hivep3*^−/−^ cells compared to C57BL/6 cells, irrespective of subset (**Fig 7c**), in agreement with higher *Cd247* transcript levels in *Hivep3*^−/−^ cells (**Sup. Table II**). By contrast, accessibility of peaks surrounding the *Ccr9* and *Drosha* genes was diminished in *Hivep3*^−/−^ cells compared to C57BL/6 cells and this was most apparent in specific iNKT cell subsets (**Fig 7c**). To associate changes in chromatin accessibility between different iNKT subsets with potential transcriptional regulators, we compared the changes in ATAC-seq signal at peaks containing known TF binding motifs between all samples using ChromVAR (**Fig 7d**). As previously shown ^21^, iNKT1 cells had the highest signal at regions containing Tbox, Runt and Ets binding motifs; peaks containing TCF, NFAT and Egr binding motifs had highest signal in iNKT2 cells, and peaks containing Rorγt binding sites had the highest signal in iNKT17 cells (**Fig 7d**). Interestingly, the analysis also demonstrated that peaks containing the NF-κB binding motif were more accessible in *Hivep3*^−/−^ cells compared with C57BL/6 cells, in all three subsets (**Fig 7d**). This is in agreement with previous reports showing that Hivep3 can directly compete with NF-κB for DNA binding ^38^. Finally, *Hivep3*^−/−^ iNKT1 cells displayed an increased in accessibility in Runt and bZIP motifs-containing peaks (**Fig 7d**), perhaps predisposing them to a more terminal iNKT1 phenotype. Altogether, these results are consistent with a role for HIVEP3 in affecting chromatin accessibility and thereby modulating the expression of certain genes. However, not all differentially express genes had corresponding changes in chromatin accessibility, suggesting that the effects of Hivep3 on iNKT cell differentiation may be linked to the activity of HIVEP3 as an adaptor protein or due to indirect, or secondary, transcriptional effects.

**Figure 7.**
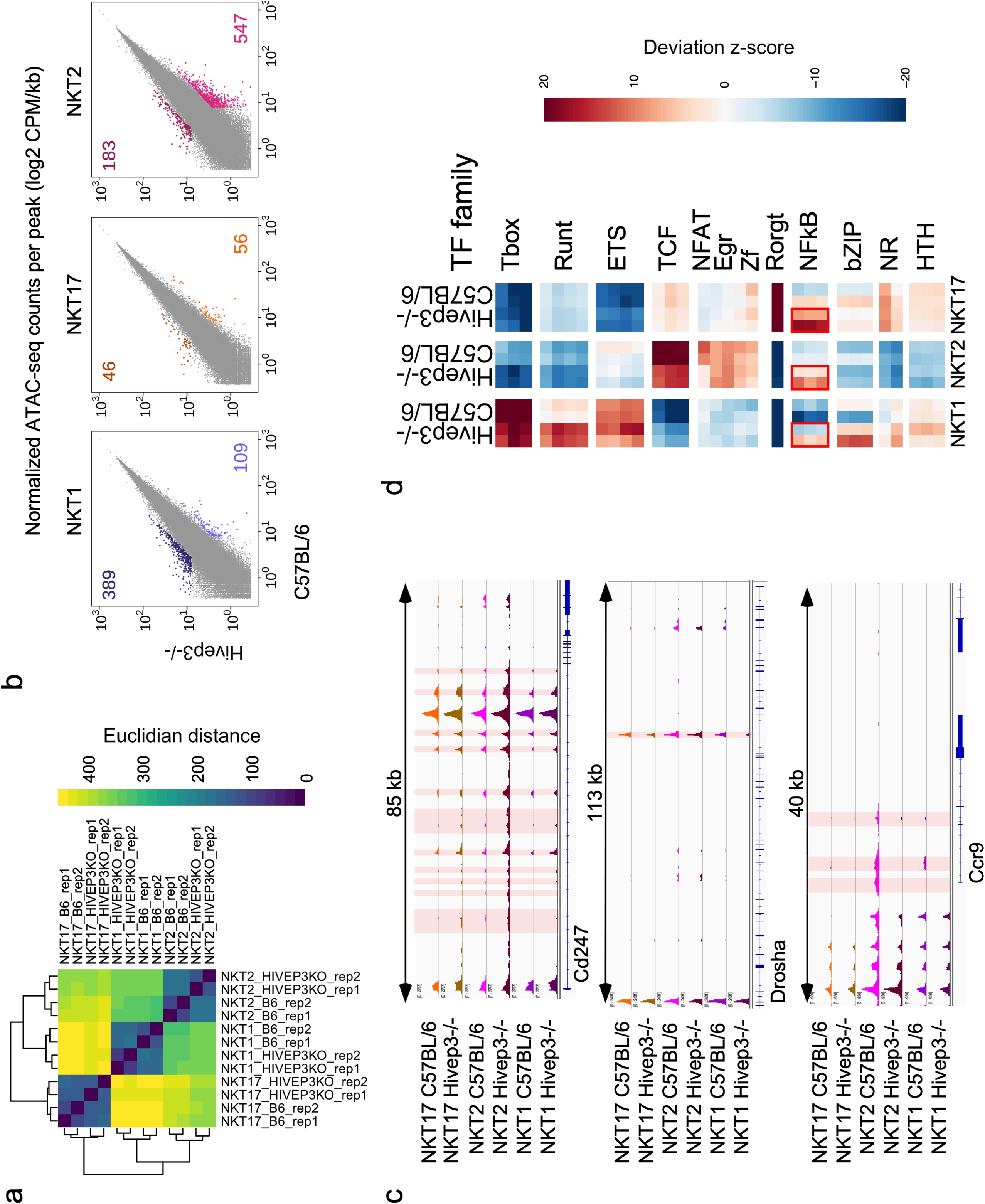
*Hivep3* deficiency impacts the chromatin landscapes of thymic iNKT cell subsets. (**a**) Euclidean distance values were computed for the iNKT1, iNKT2 and iNKT17 ATAC-seq samples from both C57BL/6 and *Hivep3*^−/−^ mice. (**b**) Scatterplots of mean ATAC-seq counts per peak comparing iNKT1, iNKT2 and iNKT17 subsets between C57BL/6 and *Hivep3*^−/−^ mice. (**c**) Mean ATAC-seq coverage at the *Cd247*, *Drosha* and *Ccr9* loci in all samples is depicted. Differentially accessible loci are highlighted in red. (**d**) Motif enrichment for iNKT1, iNKT2 and iNKT17 subsets across both strains within the differentially accessible regions is depicted as a heatmap. NF-κB binding motifs are enriched in the differentially accessible regions in the Hivep3^−/−^ samples and this is indicated with red boxes.

### PLZF^+^ innate T cells in Hivep3-deficient mice

PLZF expression is necessary and sufficient for many of the salient features that characterize T_inn_ cell function and phenotype. Because the expression of PLZF in iNKT cells was altered in absence of *Hivep3* expression (**Fig 4f**), we wondered whether *Hivep3* might also be expressed during the development of other T_inn_ cells and whether its absence would similarly affect their development. The transcriptional heterogeneity and developmental path of MAIT thymocytes was recently examined by scRNA-seq using the Chromium platform ^9^. We re-analyzed this MAIT dataset using the same parameters as for iNKT cells, corrected for batch effects and integrated the data with our sc-RNAseq iNKT datasets using the fastMNN algorithm (**Fig 8a** **and** **b**). In agreement with the emerging model that MAIT cells mirror iNKT cell development 9, both iNKT and MAIT cells distributed similarly on the UMAP and populated all clusters (**Fig 8b**). As expected, MAIT cells were found predominantly in cluster 6, which correspond to cells with a T_H_17-like transcriptome, while only a few iNKT cells populated this cluster (**Fig 8b**). Instead, a larger fraction of iNKT cells than MAIT cells occupied the clusters 7, 8, 9 10 and 11, which correspond to cells with a T_H_1-like transcriptome. These results were in good agreement with the expected frequencies for each of these subsets in both populations ^39^ and provided confidence in the integration step. In this way, we identified early CD24^+^ MAIT cells progenitors (in cluster 0), with a transcriptome akin to stage 0 iNKT cells (**Fig 8c**), as verified by the expression of *Itm2a* and *Cd24a* (**Fig 8c**). Importantly, *Hivep3* expression in MAIT cells mirrored its expression in iNKT cells, with high expression in early stage 0 cells and subsequent decline (**Fig 8c, d**). These results demonstrate that, similar to iNKT cells, *Hivep3* is also expressed early in the development of MAIT cells in the thymus. In mice, the proportion and total number of MAIT cells is much lower than for iNKT cells, making their detection more challenging.

**Figure 8.**
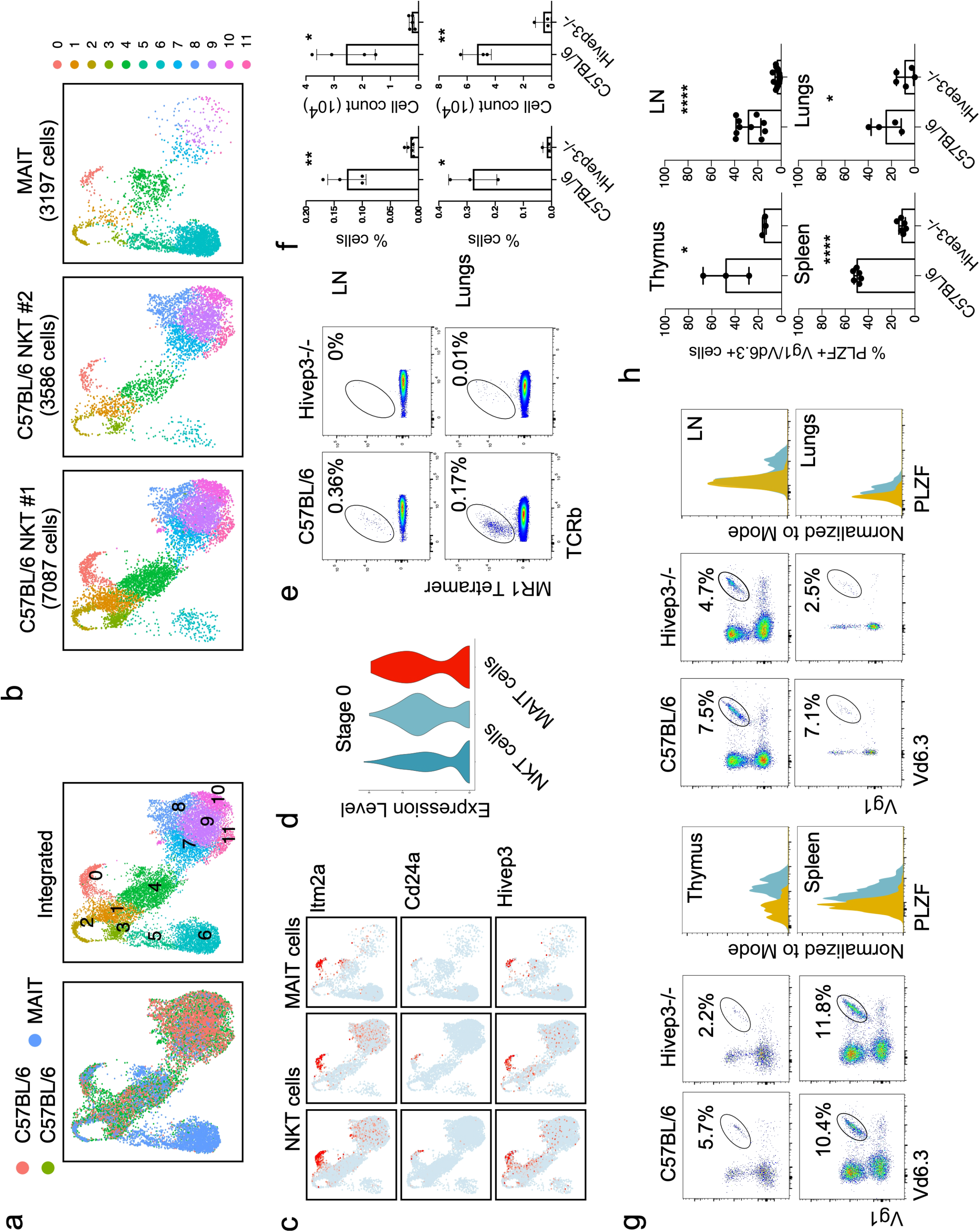
*Hivep3* deficiency universally impacts PLZF-expressing innate-like T lymphocytes. (**a**) *Left*, UMAP displaying two iNKT and one MAIT scRNA-seq sample colored by sample of origin after MNN batch-correction. *Right*, UMAP colored by inferred cluster identity for the integrated dataset. (**b**) Individual UMAP plots for each biological sample in the dataset, with the sample and number of cells passing quality control and filtering indicated above each plot. (**c**) Expression of selected genes in the iNKT and MAIT scRNA-seq samples. Each dot represents one cell and gene expression is plotted along a colorimetric gradient, with red corresponding to high expression. (**d**) Violin plots displaying aggregate expression levels of *Hivep3* in stage 0 cells for each sample. (**e**) MAIT cells in C57BL/6 and *Hivep3*^−/−^ mice were identified in the indicated tissues by staining with the MR1 tetramer and an antibody targeting TCRβ. (**f**) Quantified MAIT cell proportions and numbers in C57BL/6 and *Hivep3*^−/−^ mice in each of the indicated tissues. (**g**) Vγ1^+^ Vδ6.3^+^ γδT cells in C57BL/6 and *Hivep3*^−/−^ mice were identified in the indicated tissues by staining with antibodies targeting Vγ1 and Vδ6.3. PLZF expression in these cells from each tissue is plotted as an overlaid histogram with C57BL/6 cells colored in blue and *Hivep3*^−/−^ cells colored in yellow. (**h**) Quantified PLZF^+^ Vγ1^+^ Vδ6.3^+^ cell proportions in C57BL/6 and Hivep3^−/−^ mice in each of the indicated tissues.

Nevertheless, in our mouse colony, we can confidently detect MAIT cells in the inguinal lymph nodes and lungs using MR1 tetramers loaded with the agonist antigen 5-OP-RU (**Fig 8e**). *Hivep3*^−/−^ mice exhibited a large reduction in the proportion and numbers of MAIT cells in these organs (**Fig 8e** **and** **f**). Altogether, these results demonstrate that early in the developmental path of MAIT cells, Hivep3 is expressed and that in its absence, the proportion and numbers of MAIT cells are strongly diminished, thus mirroring the iNKT cell developmental pathway.

A subset of mouse γδT cells has also been shown to express high levels of PLZF ^40,41^. These Vγ1^+^Vδ6.3^+^ T (known as γδNKT) cells have the innate T cell capacity to rapidly secrete cytokines and chemokines upon stimulation, and this functionality is dependent upon PLZF expression ^17^. Analysis of the thymus, spleen, lymph nodes and lungs of C57BL/6 and *Hivep3*^−/−^ mice demonstrated a significant decrease in the proportion of Vγ1^+^Vδ6.3^+^ T cells amongst total γδT cells in all organs analyzed (**Fig 8g**). Importantly, we did not detect any expression of PLZF in the remaining Vγ1^+^Vδ6.3^+^ T cells from *Hivep3*^−/−^ mice, in contrast to what is observed in the comparable population found in C57BL/6 mice (**Fig 8g** **and** **h**). Furthermore, in the course of our analysis we observed a population of αβ T cells in the inguinal lymph nodes that were neither iNKT nor MAIT cells, and yet co-expressed the PLZF and Rorγt transcription factors (**Sup. Fig 7**). These cells, which are predominantly of the CD4^−^ CD8^−^ phenotype, were also present in the inguinal lymph nodes of CD1d1d2^−/−^ mice, indicating that they do not correspond to type II NKT cells. Although the exact nature of this population remains to be determined, these cells were nevertheless largely absent from the inguinal lymph nodes of *Hivep3*^−/−^ mice (**Sup. Fig 7**).

Interestingly, innate lymphoid cells (ILCs), which also require PLZF expression early in their development ^42^, were detected in normal proportions and numbers in the lungs of Hivep3-deficient mice (**Sup. Fig 8a, b**), suggesting that requirement for *Hivep3* expression does not extend to all PLZF-expressing immune cells. Additionally, we examined whether absence of Hivep3 expression impacts the development of Foxp3^+^ regulatory T cells ^43^ and the precursors of intraepithelial lymphocytes ^44^, two PLZF^−^ T cell populations known to require “agonist” selection ^45^, just as T_inn_ cells do. Both thymic populations were found at comparable frequencies and numbers between C57BL/6 and *Hivep3*^−/−^ mice (**Sup. Fig 8c, d, e** **and** **f**). Altogether, our results demonstrate that *Hivep3*-deficiency specifically affects T_inn_ cells that express PLZF.

## Discussion

Exhibiting characteristics of both innate and adaptive immunity, innate-like T lymphocytes such as iNKT cells, MAIT cells, and subsets of γδT cells, have emerged as key players in the control of immunity and tissue homeostasis ^46–49^. In striking contrast to T_conv_ cells, which exit the thymus in a “naïve” state requiring an orchestrated process of priming and clonal expansion over several days to become fully functional effector cells, T_inn_ cells can acquire most of their functions, such as the capacity to secrete multiple cytokines upon primary stimulation, a memory/activated phenotype (CD44^hi^ CD62L^−^) associated with their distribution within tissues and cytotoxic potential, during their thymic development. These unique innate-like features are directed, at least in part, by the expression of the *Zbtb16*-encoded transcription factor PLZF, which commonly regulates the acquisition of T-helper effector programs during the development of innate-like lymphocyte lineages ^50,51^. However, a complete mechanistic understanding of T_inn_ differentiation and function remains elusive. Detailed knowledge of the gene expression changes driving progression from one cellular state to the next is lacking. To this end, we used scRNA-seq to investigate the transcriptional landscape of iNKT cell maturation and fate decisions under steady-state conditions. Our results outline the transcriptional signature of the previously identified stage 0, iNKT1, iNKT2 and iNKT17 cells, and further expose heterogeneity amongst iNKT1 cells. Surprisingly, despite the large number of cells analyzed (>10,000) and our enrichment for immature CD44^low^ cells, we did not detect cells with transcriptional signatures indicative of the presence of iNKT10 (ref ^52^) or iNKT_FH_ subsets ^53^ within the thymus, suggesting that such subsets likely arise as a consequence of further differentiation within peripheral organs. iNKT17 cells were transcriptionally homogenous, forming only one cluster in our analysis. However, iNKT2 and especially iNKT1 cells were more heterogenous and contained transcriptionally diverse cells. *Zbtb16* transcripts were first detected in a subset of stage 0 cells, immediately adjacent on the UMAP to three clusters of cells defined by their S, S/G2 and G2 cell cycle scores, respectively. These results are in agreement with the intra-thymic proliferative expansion that occurs post-stage 0 iNKT cells ^5,28^. Interestingly, these cells were enriched for a transcriptional signature previously ascribed to intermediary precursor cells, iNKTp ^27^, and were enriched for *Ccr7*, *Ccr9* and *Il13*-encoding transcripts. This IL-13-expressing intermediate population was previously shown to give rise to all mature iNKT1, iNKT2, and iNKT17 subsets ^37^. Cells with an otherwise similar transcriptional profile to iNKT2 cells but lacking expression of cell-cycle related transcripts were also observed, representing *bona-fide* iNKT2 cells (*Zbtb16*^+^, *Icos*^+^, *Izumo1r*^+^, *Il6ra*^+^, *Il4*^+^ but *Il13*^−^). Surprisingly, within this cluster of iNKT2 cells, we detected cells in “bridge” regions of the UMAP with transcriptional signatures straddling those of iNKT1 and iNKT17 cells. These results suggest that iNKT2 cells might not solely represent a terminally differentiated subset of iNKT cells as previously thought ^8^, but instead also include cells undergoing an ongoing process of differentiation towards an iNKT17 or iNKT1 transcriptional profile. Such an interpretation is supported by analysis of developmental trajectories yet appears to conflict with previous intra-thymic transfer experiments showing that IL-4-secreting iNKT2 cells maintained their phenotype 4 days post-transfer ^8^. It is possible that in these experiments, further differentiation of iNKT2 cells into iNKT1 or iNKT17 cells was not observed because this process requires a longer period of time than the one analyzed and/or that the regions of the thymus into which the cells were injected were not conducive to further differentiation. Alternatively, pseudo-time inference analyses rely on the presence of a sufficient number of cells with transitional transcriptional profiles to place cells along a pseudo-time trajectory. Despite the analyses of over 10,000 cells in our study, it remains possible that our current dataset did not capture enough transitional cells, thereby affecting the interpretations of developmental trajectories. Further sc-RNAseq experiments aimed at analyzing the transcriptional diversity of IL4-producing iNKT2 cells and the temporal development of iNKT cells, associated with fate-mapping and *in vivo* transfer experiments, should help further refine our understanding of the developmental path undertaken by thymic iNKT cells. Unexpectedly, we also observed transcriptional diversity amongst iNKT1 cells, suggesting a progressive maturation process in which, concomitant with the detection of transcripts encoding Ly6C and killer cell lectin type receptors, cells gained expression of *Ifng* transcripts and cytotoxic molecules (*Gzmb*, *Gzma*). A clear cluster of iNKT1 cells with type I interferon gene response signature was also identified. In future experiments, it will be interesting to evaluate whether type I interferon signaling in the thymus plays a role in the functional maturation of iNKT1 cells.

Our dataset captured a large number (565 cells) of stage 0 iNKT cells, allowing for a detailed analysis of the early transcriptional events that characterize the commitment towards the innate effector fate. We identified expression of several TF indicative of TCR signaling in stage 0 cells, several of which had previously been implicated in the development of iNKT cells. However, we also noted high and transient expression of *Hivep3* transcripts in stage 0 cells. HIVEP3 belongs to the ZAS family of proteins ^15^. Genes in this family encode for large proteins with two separate Cys2His2 zinc finger pairs that independently bind to specific DNA sequences. HIVEP3 was initially cloned from a mouse thymocyte cDNA expression library using the consensus signal sequence of somatic V(D)J recombination (RSS) as a ligand ^54^ and was later shown to have dual DNA specificity for the kB motif ^55^. HIVEP3 was also shown to act as an adaptor protein. By binding to TRAF2 (ref ^13^), HIVEP3 can inhibit both NF-κB and c-Jun NH2-terminal kinase (JNK)-mediated responses, while by physically interacting with c-Jun it serves as a co-activator of AP-1-dependent Il-2 gene transcription ^14^. Here, we demonstrate that HIVEP3 serves as a gatekeeper universally controlling the development of PLZF^+^ innate T cells, including iNKT, MAIT and γδNKT cells. In iNKT cells, the absence of HIVEP3 affected the expression of multiple genes. For some of these genes, HIVEP3 probably disturbed their expression by modifying chromatin accessibility of regulatory regions. Indeed, our ATAC-seq analysis of iNKT cell subsets clearly demonstrated differential chromatin accessibility surrounding certain gene loci that correlated with gene expression. Furthermore, in agreement with previous results showing that Hivep3 can directly compete with NF-κB for binding to the κB motif, we also found increased accessibility in regions of the genome with a κB motif when HIVEP3 was absent.

Future chromatin immunoprecipitation experiments of HIVEP3 should reveal whether such effects are mediated directly by HIVEP3 bound to the DNA or whether the effects are indirect. We also observed changes in gene expression that were not associated with any apparent change in chromatin accessibility. In such cases, modulation of gene expression might have been mediated through the adaptor activity of HIVEP3. The mechanistic and possible binding partners of HIVEP3 in iNKT cells remain to be explored. Moreover, our results revealed a near absence of transcripts encoding the microRNA processing enzyme Drosha in *Hivep3*-deficient iNKT cells. T_inn_ cells are particularly sensitive to disruption of miRNA function, both globally and on the individual miRNA level ^56–59^. We speculate that with limited Drosha expression, miRNAs processing might have been affected and contributed to the dysregulation of iNKT development. In support of this hypothesis, we observed that a large fraction of DEGs showed higher levels of expression in *Hivep3*-deficient cells compared to C57BL/6 cells, perhaps reflecting a deficiency in post-transcriptional regulation of gene expression. Future experiments analyzing miRNAs expression in *Hivep3*-deficient iNKT cells will explore this possibility. The class IIa histone deacetylase HDAC7 is highly expressed in double-positive thymocytes where it resides in the nucleus ^60^. Upon thymic selection, HDAC7 is exported out of the nucleus in a TCR-dependent manner, which thereby affects the transcriptional program of developing T cells ^61^. We observed an increase in *Hdac7*-transcript levels in stage 0 iNKT cells of *Hivep3*^−/−^ mice compared to their wildtype counterparts. Although the localization of HDAC7 in these cells is currently unknown, it is noteworthy that transgenic mice with a T-cell specific and transient expression of a mutant HDAC7 lacking the phosphorylation sites required for TCR-dependent nuclear export (HDAC7-ΔP), exhibit a dramatic loss of iNKT cells due to developmental defects. In these mice, the intra-thymic proliferation of iNKT cells is prevented and expression of PLZF is transcriptionally repressed ^34^. HDAC7 was also shown to physically interact with PLZF and affects its transcriptional activity. These results resonate with our observations in *Hivep3*^−/−^ mice showing a block in iNKT cell development with reduced thymic expansion and decreased PLZF expression. Interestingly, modulation of PLZF protein expression early in the course of iNKT development was previously shown to result in significant reduction in iNKT cell numbers ^62,63^, providing another potential mechanism by which HIVEP3 might control the development of iNKT cells.

Reinforcing the prevailing idea that MAIT cells follow a developmental program analogous to iNKT cells ^9^, we found that *Hivep3*-deficient mice had a profound defect in MAIT cells and other populations of T_inn_ cells. Interestingly, innate lymphoid cells that also require PLZF for their development were not affected by the absence of *Hivep3* expression, suggesting that HIVEP3 requirement does not extend to all PLZF^+^ immune cells but may be restricted to TCR expressing cells. This would be consistent with the induction of Hivep3 expression in T cells upon TCR engagement ^14^, and upon positive selection during their development ^64^. However, we did not detect obvious deficiencies in T_conv_ or in other “agonist” selected T cells, such as Tregs and IELps in *Hivep3*^−/−^ mice. Like iNKT and MAIT cells, these agonist selected cells require strong agonist TCR signaling during selection. However, they do not need PLZF expression for proper development and are not selected by MHC-expressing DP thymocytes ^45^. Overall, these results point toward the intriguing possibility that HIVEP3 activity might be uniquely required by T cells that, in addition to strong TCR signaling during selection, also require homotypic interactions between signaling lymphocyte activation molecule family (SLAMF) receptors, SLAM and Ly108. Perhaps by providing such unique co-signals during positive selection, it distinctively affects the many potential protein modification sites on HIVEP3. For example, the phosphorylation status of HIVEP3 was previously shown to modulate its DNA-binding affinity by 9-fold ^65^. Altogether, these findings identify a new and key regulator of the naïve versus PLZF^+^ innate T lymphocytes developmental choice.

## Material and Methods

### Mice

The *Hivep3*^−/−^ mice backcrossed to the C57BL/6 background have been described previously and were graciously provided by Dr. Laurie Glimcher. C57BL/6 and CD45.1 congenic C57BL/6 mice were purchased from Jackson Laboratories. All mice were used between 8 to 10 weeks and were age matched for each experiment. All mice were raised in a specific pathogen-free environment at the Office of Laboratory Animal Research at the University of Colorado Anschutz Medical campus. All animal procedures were approved by the UCD (00065) Institutional Animal Care and Use Committees and were carried out in accordance with the approved guidelines.

### Thymocyte isolation and flow cytometry

Single cell suspensions were prepared from the thymus by manual disruption using a syringe plunger. PBS57-CD1d1 and 5-OP-RU-MR1 tetramers were obtained from the National Institutes of Health Tetramer Core Facility. The complete list of surface antibodies used is as follows: from BioLegend – CD3 (clone 17A2; Catalog #100204, Catalog #100218), CD4 (clone GK1.5; Catalog #100447), CD4 (clone RM4-5; Catalog #100557), CD19 (clone 6D5; Catalog #115521, Catalog #115528), CD44 (clone IM7; Catalog #103044), CD45.1 (clone A20; Catalog #110714), CD45.2 (clone 104; Catalog #109828), CD69 (clone H1.2F3; Catalog #104530), CD81 (clone Eat-2; Catalog #104913), ICOS (clone 7E.17G9; Catalog #117406, Catalog #117424), NK1.1 (clone PK136; Catalog #108730, Catalog #108736), PD-1 (clone 29F.1A12; Catalog #135206), TCRβ (clone H57-597; Catalog #109220, Catalog #109230) and Vγ1 (clone 2.11; Catalog #141104, Catalog #141108); from BD Biosciences – CD8α (clone 53-6.7; Catalog #563786), CD24 (clone M1/69; Catalog #563545) and Vδ6.3 (clone 8F4H7B7; Catalog #555321).

After surface antibody staining, the cells were fixed and permeabilized using the FoxP3 fixation/permeabilization kit (Thermo Fisher Scientific). Fixed and permeabilized cells were incubated with a combination of the following antibodies targeting intracellular proteins: from Abcam – Lef-1 (clone EPR2029Y; Catalog #ab137872); from BioLegend – Bcl-2 (clone BCL/10C4; Catalog #633512), Helios (clone 22F6; Catalog #137222), IFNγ (clone XMG1.2; Catalog #505826, Catalog #505830), IL-4 (clone 11B11; Catalog #504104), PLZF (clone 9E12; Catalog #145808) and T-bet (clone 4B10; Catalog #644816, Catalog #644824); from BD Biosciences – RORγt (clone Q31-378; Catalog #562684); from Miltenyi Biotec – Tox (clone REA473; Catalog #130-118-335); from Thermo Fisher Scientific – Egr2 (clone erongr2; Catalog #12-6691-82, Catalog #17-6691-82, Catalog #25-6691-82), Ki-67 (clone SolA15; Catalog #25-5698-82) and PLZF (clone Mags.21F7; Catalog #53-9320-82, Catalog #25-9322-82).

The stained cells were then analyzed on a BD LSRFortessa (BD Biosciences) or Cytek Aurora (Cytek) and data were processed with FlowJo software vX (TreeStar).

### Enrichment of CD1d reactive thymocytes

Thymocytes were enriched for PBS57-CD1d reactive cells by incubating thymocyte cell suspensions with PE or APC conjugated PBS57-CD1d tetramers for 45 minutes at 4°C, then incubated with anti-PE or anti-APC magnetic microbeads (Miltenyi Biotec) for 15 minutes at 4°C, followed by separation by using an autoMACS Pro Separator (Miltenyi Biotec) according to manufacturer’s instructions. Subsequently, cells in the positive fraction were first stained for surface markers followed by staining for intracellular markers before being subjected to flow cytometric acquisition.

### Lung single cell suspensions

To prepare single-cell suspensions lungs were finely chopped with scissors in a 48 well plate and treated with 3 mg ml^-1^ collagenase III (Worthington, Lakewood, NJ), 5 μg ml^-1^ DNAse, and complete media (RPMI containing 10% fetal calf serum and enriched supplements) for 60 min at 37°C with gentle pipetting at 20 min intervals. Cells were then filtered through a 70 μm cell strainer and washed with complete media.

Approximately 2 × 10^6^ cells were filtered (40 μm) and used for flow cytometric analysis.

### Liver single cell suspensions

Hepatic leukocytes were isolated by cutting individual livers into small pieces and gently pressed through a 70 μm filter placed on top of a 50 ml falcon tube and resuspended in FACS buffer (PBS, 0.5% BSA, 0.5 mM EDTA, 1% Azide). The cells were washed twice in ice-cold FACS buffer and spun through 33.8% Percoll (Amersham Pharmacia Biotech) for 12 min at 2000 rpm, room temperature. Red blood cells were removed by resuspending the pellet in red blood cell lysis buffer for 5 min at room temperature and washed with complete media. Cells were resuspended in FACS buffer and filtered through a 40 μm cell strainer.

### Bone marrow chimeras

Bone marrow cells from congenic C57BL/6 mice (CD45.1) and *Hivep3*^−/−^ (CD45.2) mice were harvested and depleted of CD90-expressing cells using magnetic beads. Bone marrow cells from CD45.1 mice were mixed at 1:1 ratio with CD45.2 *Hivep3*^−/−^ mice. 5 ×10^6^ cells were injected intravenously into lethally-irradiated recipient mice (1000 rads). Eight to ten weeks after reconstitution, chimeric mice were sacrificed and thymocytes were analyzed by flow cytometry.

### BrdU incorporation assay

Mice received an intraperitoneal injection of BrdU (BrdUrd; 2 mg in 200 μL PBS; BD Biosciences) or PBS only at 0, 24, and 48 h before being sacrificed for analysis at 72h. To detect the incorporation of BrdU, the BrdU Flow Kit (BD Biosciences, Catalog #552598) was used according to the manufacturer’s protocol.

### *In vivo* stimulation of iNKT cells

Mice received 2 μg of αGC (Alexis Biochemicals) or vehicle by intravenous injection. Ninety minutes after injection, organs were collected and processed. Cells were first surface stained, fixed and subsequently stained intracellularly for lL-4 and IFNγ.

### scRNA-seq data processing

The quality of sequencing reads was evaluated using FastQC and MultiQC. Cell Ranger v3.1.0 was used to align the sequencing reads (FASTQ) to the mm10 mouse transcriptome and quantify the expression of transcripts in each cell. This pipeline produced a gene expression matrix for each sample, which records the number of UMIs for each gene associated with each cell barcode. Unless otherwise stated, all downstream analyses were implemented using R v3.6.1 and the package Seurat v3.1.4 (ref ^66^). Low-quality cells were filtered using the cutoffs nFeature_RNA >= 500 & nFeature_RNA < 4100 & nCount_RNA >=1000 & percent.mt < 6. The NormalizeData function was performed using default parameters to remove the differences in sequencing depth across cells. Dimension reduction was performed at three stages of the analysis: the selection of variable genes, PCA, and uniform manifold approximation and projection (UMAP) ^67^. The FindVariableGenes function was applied to select highly variable genes covering most biological information contained in the whole transcriptome. To remove batch-effect from the PCA subspaces based on the correct cell alignment, we used fastMNN ^18^ to detect mutual nearest neighbors (MNN) of cells in different batches, and then used the MNN to correct the values in each PCA subspace. The variable genes were used for MNN as input to perform the RunUMAP function to obtain bidimensional coordinates for each cell. We determined the k-nearest neighbors of each cell using the FindNeighbors function and used this knn graph to construct the SNN graph by calculating the neighborhood overlap (Jaccard index) between every cell and its k.param nearest neighbors. Finally, we used the FindClusters function (resolution 0.8) to cluster cells using the Louvain algorithm based on the same PCs as RunUMAP function.

### Identification of differentially expressed genes

We identified cluster-enriched genes by using the FindAllMarkers function in Seurat and the Wilcoxon-Rank sum test. This function identified differentially expressed genes for each cluster by comparing the gene expression for cells belonging to a cluster versus cells belonging to all other clusters. Only those genes that passed an adjusted *p* value (Benjamini-Hochberg) cutoff of 0.05 were included in the downstream analyses. Gene ontology (GO) analysis was performed by using the R package clusterProfiler (v3.0.4) ^68^.

### Developmental trajectory inference

Two independent trajectory packages (Slingshot and Monocle v3) were used to order the cells in pseudotime. To run Slingshot, we first converted the Seurat object into a SingleCellExperiment object. We then implemented the Slingshot algorithm in a semi-supervised manner by providing this object as input to the slingshot function and specifying the start cluster as cluster 0 (start.clus = ‘0’). This function then determines the global lineage structure using cluster-based minimum spanning tree and subsequently fitting simultaneous principal pseudotime curves to describe each determined lineage. Lastly, we plotted the lineage curves on the UMAP plot generated by the Seurat package. We then fit a general additive model (GAM) for each curve to identify the genes whose expressions vary along the curve. For the Monocle 3 pipeline, we followed the recommended workflow to create a cell_data_set object with our cells. Since the UMAP coordinates for the cells provided by Monocle 3 were not equivalent to the coordinates provided by Seurat, we replaced the UMAP coordinates for each cell in the cell_data_set object with those from the Seurat object. We then ordered the cells using the order_cells function and choosing cluster 0 cells as the root node.

The trajectories were then plotted on the UMAP using the plot_cells function and setting the color_cells_by argument to ‘pseudotime’.

### ATAC-Seq

ATAC-seq was performed according to the Omni-ATAC protocol previously described by Corces et al. ^69^. 5,000 to 20,000 cells from each iNKT subset (iNKT1 cells (CD122^+^, ICOS^−^), iNKT2 cells (CD122^−^, ICOS^+^, Izumo1r^+^, CD138^−^) or iNKT17 cells (CD122^−^, ICOS^+^, Izumo1r^−^, CD138^+^)) from the pooled thymi of C57BL/6 or Hivep3^−/−^ mice were sorted individually and then pooled for a total of 50,000-100,000 cells per transposition (). Cells were then placed in 50 mL of cold lysis buffer (10 mM TrisHCl, pH 7.4, 10 mM NaCl, 3 mM MgCl2, 0.1% (v/v) Molecular biology-grade NP-40, 0.1% (v/v) Tween-20, 0.01% (v/v) Digitonin).

Permeabilized cells were pelleted for 10 minutes at 500xg and resuspended in 50 μL of transposition mix (25 μL 2x TD buffer (Illumina), 16.5 μL 1X PBS, 0.5 μL 10% (v/v) Tween-20, 0.5 μL 1% (v/v) Digitonin, 2.5 μL Tn5 transposase (Illumina), 5 μL nuclease-free water). The volume of lysis buffer/transposition mix was scaled up to 100 μL for 100,000 cells. The transposase reaction was conducted for 30 min at 37 °C with shaking at 1,000 rpm. Cells were then subjected to a second sort to separate the different iNKT cell subsets using the same markers used in the first sort. While the fluorescence corresponding to the CD122 and Izumo1r markers was quenched by the transposition reaction, other markers were unaffected, allowing for the sorting of each of the iNKT cell subsets on the basis of ICOS and CD138 expression (iNKT1 (ICOS^−^ CD138^−^), iNKT2 (ICOS^+^, CD138^−^) and iNKT17 (ICOS^+^ CD138^+^)). Chromatin accessibility surrounding subset-defining genes (i.e. *Tbx21*, *Rorc* and *Slamf6*) is in accordance with the expected pattern of each iNKT subset (**Sup. Fig 6**). DNA was purified using the DNA Clean and Concentrator-5 kit (Zymo Research) and barcoding was performed using Illumina compatible index primers designed according to Buenrostro et al. ^70^ (Integrated DNA Technologies). PCR was conducted for 11–12 cycles. Library purification and size selection was carried out using 1.8X AMPureXP beads (Beckman Coulter). Libraries were quantified and size distribution was assessed using the High Sensitivity D1000 ScreenTape System (Agilent).

Paired-end sequencing was performed on a NovaSeq (Illumina) with 150 cycles for each read. Raw data from the sequencer was demultiplexed and FASTQ files were generated using bcl2fastq conversion software (Illumina).

### ATAC-Seq Analysis

Sequencing reads in FASTQ format were trimmed to 50 bp using Trimmomatic and mapped to mouse genome (mm10) using the ENCODE ATAC-seq pipeline, with default parameters, except mapped reads were subsampled to a maximum of 50 million reads for peak calling. NarrowPeak files from individual replicates of all conditions were merged into a global set of peaks, excluding those on the Y chromosome or overlapped the ENCODE blacklist, and condensed to non-overlapping regions with a uniform size of 500 bp using chromVAR ^71^. The number of transposase insertions within each region was computed for each replicate using chromVAR and these raw ATAC-seq counts per peak for all replicates were normalized using voom. Pairwise contrasts were performed with limma and differentially accessible regions were filtered based on an FDR adjusted p-value of less than 0.1 and an estimated fold-change of at least 3. We computed the ATAC-seq density (number of transposase insertion sites per kilobase) and accessible regions were defined as those with a mean of 5 normalized insertions per kilobase. We associated transcription factors binding motifs from the HOMER database by determining the enrichment of motifs in groups of peaks with HOMER and comparing the variability in ATAC-seq signal with chromVAR. For visualization, genomic coverage for individual replicates were computed on 10 bp windows with MEDIPS using full fragments captured by ATAC-seq and used to generate average coverage with the Java Genomics Toolkit for each group.

### Statistical analysis

Groups were compared with two-tailed Student’s *t*-tests or one-way analysis of variance (ANOVA) with Prism 8 (Graphpad Software, Inc.). \**P* < 0.05, \*\**P* < 0.01, \*\*\**P* < 0.001 and \*\*\*\**P* < 0.0001.

## Data availability

Data that support the findings of this study have been deposited in NCBI GEO with the accession code xxx.

## Acknowledgments

We thank members of our laboratories for thoughtful discussions and critical comments on the manuscript; Jennifer Matsuda and Leslie Berg for critical comments and support; the Flow Core and the University of Colorado flow cytometry shared resource facility for assistance with cell sorting; the Genomics and Microarray core at the University of Colorado Anschutz Medical campus for scRNA-Seq and ATAC-seq and the National Institutes of Health core facility for CD1d and MR1-tetramers. This work was supported by National Institutes of Health Grants AI135339 and AI130198 (to L.G.); The Cancer Center Support Grant P30CA046934.

## Author contributions

L. G. and S. H. K. wrote the manuscript and designed figures. L. G., S. H. K., J. Z. and J. SB. designed experiments. S. H. K., J. Z., M. J. M-F., and L. L. conducted experiments and acquired data. S. H. K., J. Z., M. J. M-F., L. L., T. B., J. SB. and L. G. analyzed and interpreted data. L. G., S. H. K., and J. SB edited the manuscript. All authors approved the final manuscript.

## Competing Financial Interests

The authors declare no competing financial interests.

---

**Supplementary Table I.** Differentially upregulated genes per cluster in C57BL/6 thymic iNKT cells.

**Supplementary Table II.** List of differentially regulated genes in C57BL/6 vs *Hivep3^−/−^* iNKT cells per cluster.

**Supplementary Figure 1.**
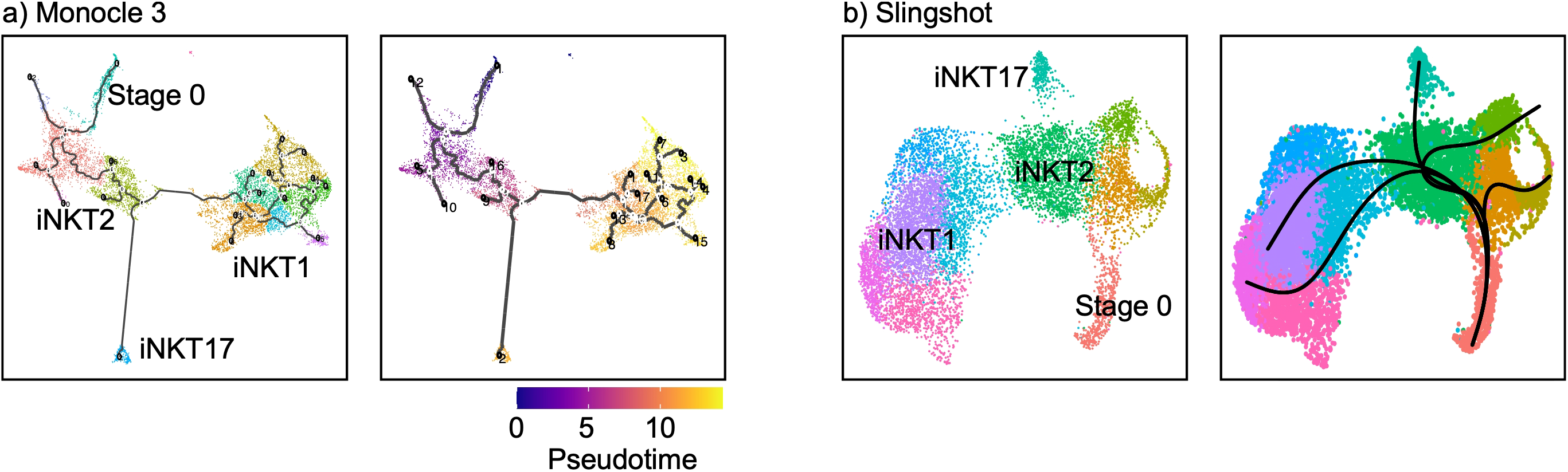
Developmental trajectories of thymic iNKT cells. (**a**) Pseudotemporal trajectory was computed using the Monocle3 trajectory inference R package and displayed on a UMAP plot. Cells are colored by both cluster identity and along their computed pseudotime values. (**b**) Pseudotemporal trajectories computed using the Slingshot trajectory inference R package on a UMAP plot. Cells in cluster 0 were provided as the root node for both softwares since those cells represent the earliest iNKT cell precursors. The softwares then determine the differentiation paths that the stage 0 cells could take based on gene expression.

**Supplementary Figure 2.**
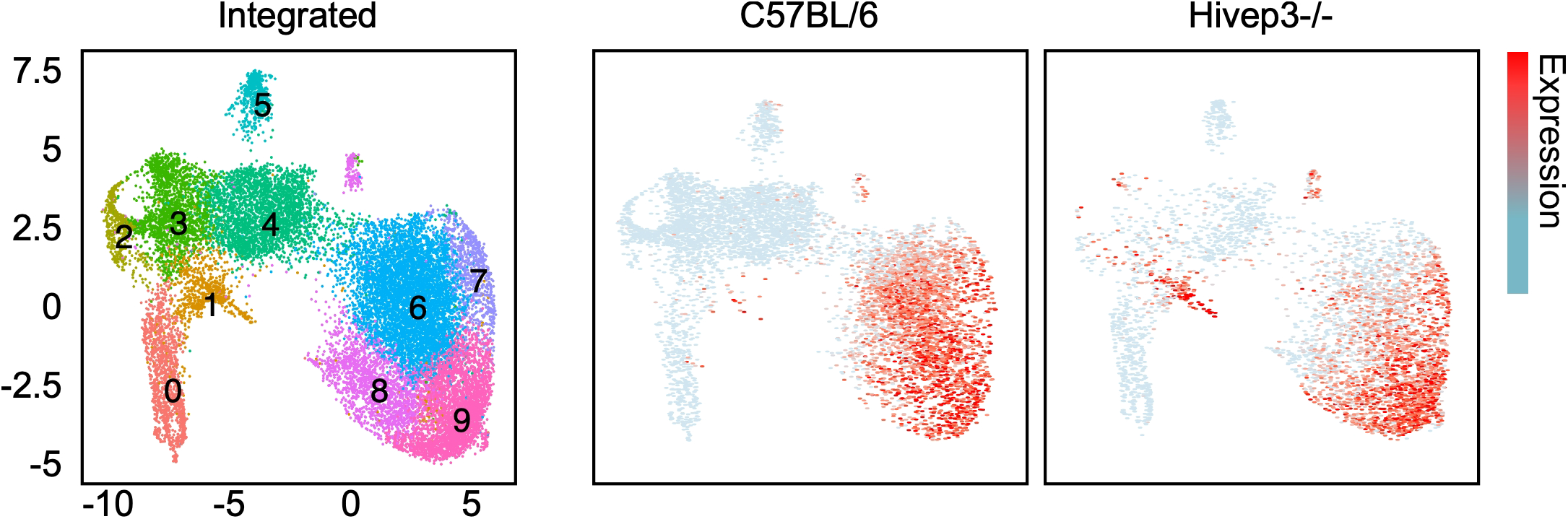
*Hivep3*^−/−^ cluster 1 cells possess a mature iNKT1 gene signature. Expression of selected iNKT1 gene signature (*Cd7*, *Dusp1*, *Ms4a4b*, *Il2rb*, *Fcer1g*, *Klhdc2*, *Klra1*, *Klf2*, *Klrb1a*, *Cd160* and *S100a4*) in the C57BL/6 and Hivep3^−/−^ single cell RNA-seq samples. Each dot represents one cell and gene expression is plotted along a colorimetric gradient, with red corresponding to high expression.

**Supplementary Figure 3.**
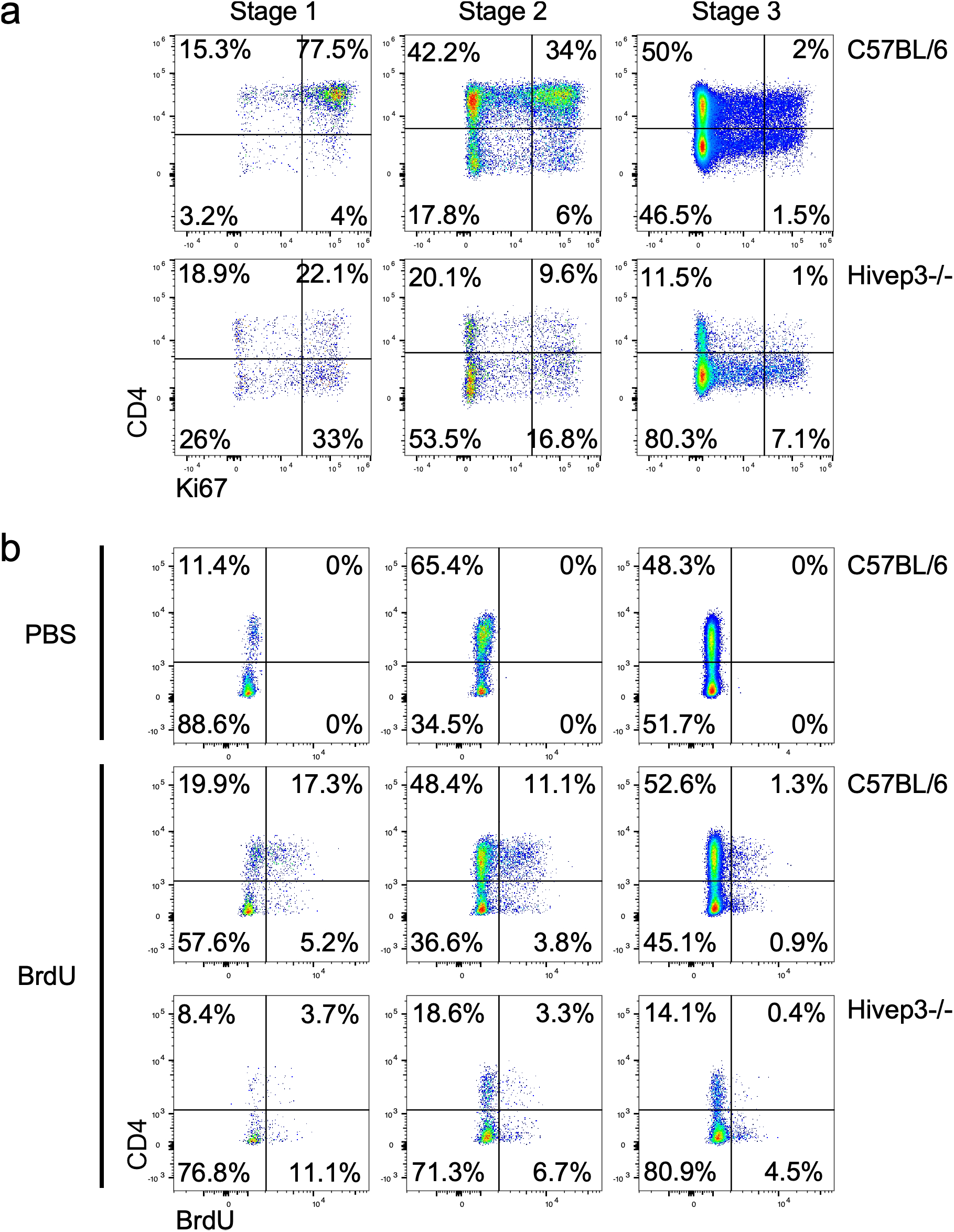
*Hivep3*^−/−^ iNKT cells display a reduced proliferative potential. **(a)** Representative flow cytometry plots depicting the expression of CD4 and Ki67 on the three developmental stages of iNKT cells (Stage 1, CD44^−^NK1.1^−^, stage 2, CD44^+^NK1.1^−^ and stage 3, CD44^+^NK1.1^+^) in the thymi of 8-week-old C57BL/6 and *Hivep3*^−/−^ mice. (**b**) Representative flow cytometry plots depicting the expression of CD4 and incorporation of BrdU by the three developmental stages of iNKT cells in the thymi of 8-week-old C57BL/6 and *Hivep3*^−/−^ mice (data representative of n = 3).

**Supplementary Figure 4.**
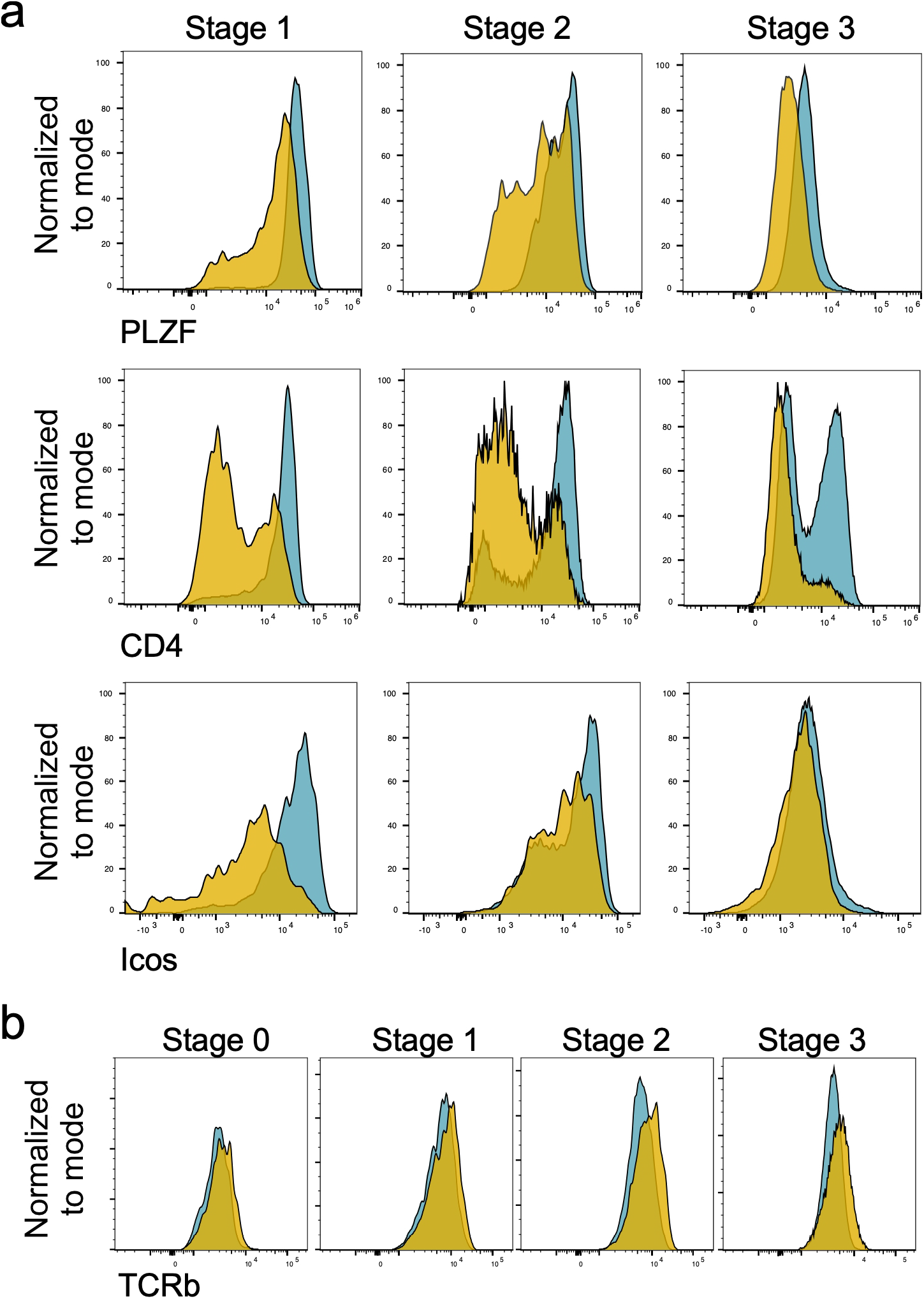
Characterizing protein expression of *Hivep3*^−/−^ iNKT cells. (**a**) Representative flow cytometry plots depicting the expression of PLZF, CD4 and ICOS on the three developmental stages of iNKT cells in the thymi of 8-week-old C57BL/6 and *Hivep3*^−/−^ mice. (**b**) Representative flow cytometry plots depicting the levels of expression of TCRβ by the four developmental stages of iNKT cells (Stage 0, CD24^+^CD44^−^NK1.1^−^, Stage 1, CD44^−^NK1.1^−^, stage 2, CD44^+^NK1.1^−^ and stage 3, CD44^+^NK1.1^+^) in the thymi of 8-week-old C57BL/6 and *Hivep3*^−/−^ mice (data representative of n = 3).

**Supplementary Figure 5.**
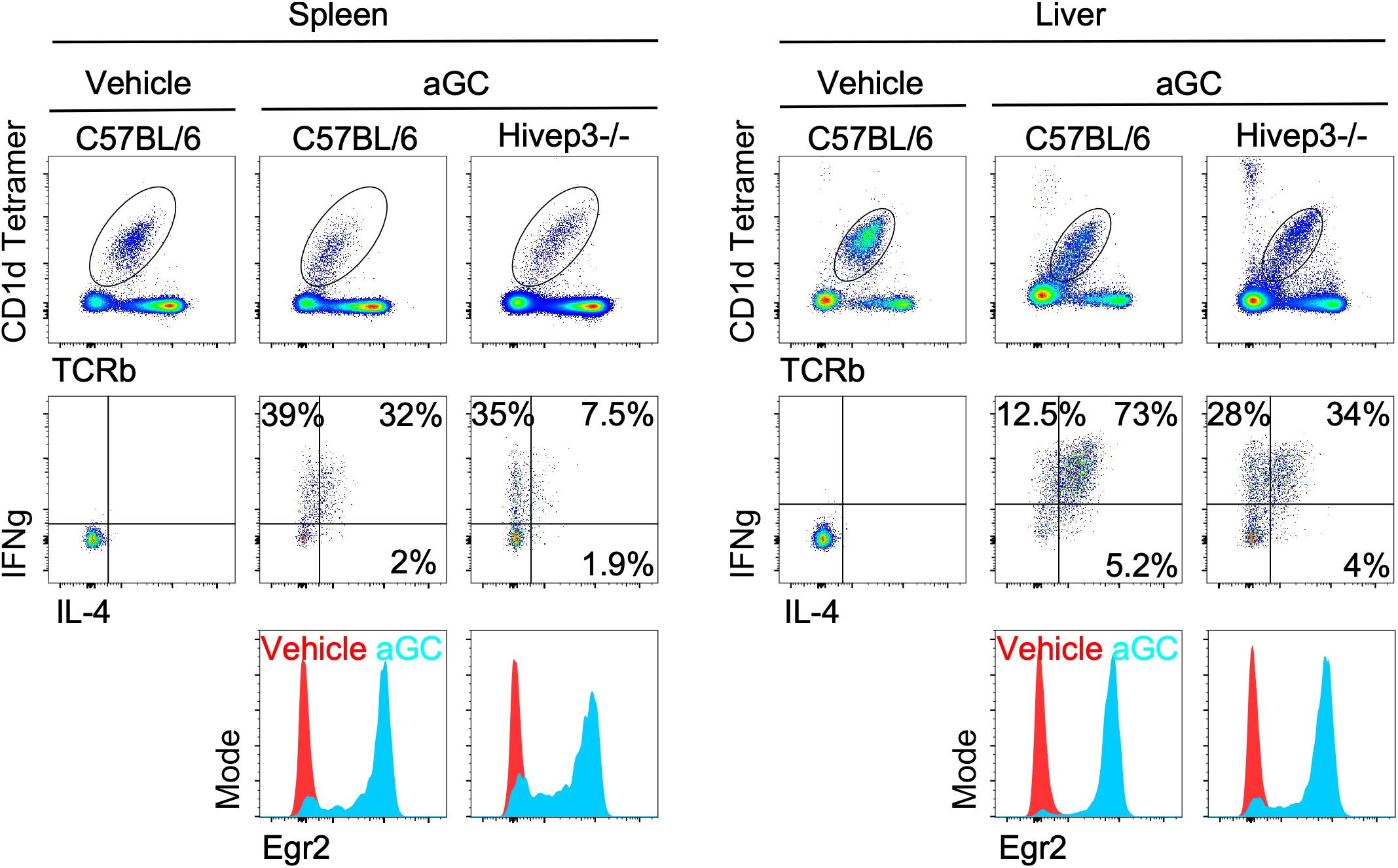
Peripheral iNKT cells in *Hivep3*^−/−^ mice are poorly responsive. iNKT cells from C57BL/6 and *Hivep3*^−/−^ mice were isolated from the spleen and liver 90 minutes following i.p. administration of the agonist lipid αGC and stained intracellularly for IFNγ and IL-4. The cells were also stained for the transcription factor Egr2 (data representative of n = 3).

**Supplementary Figure 6.**
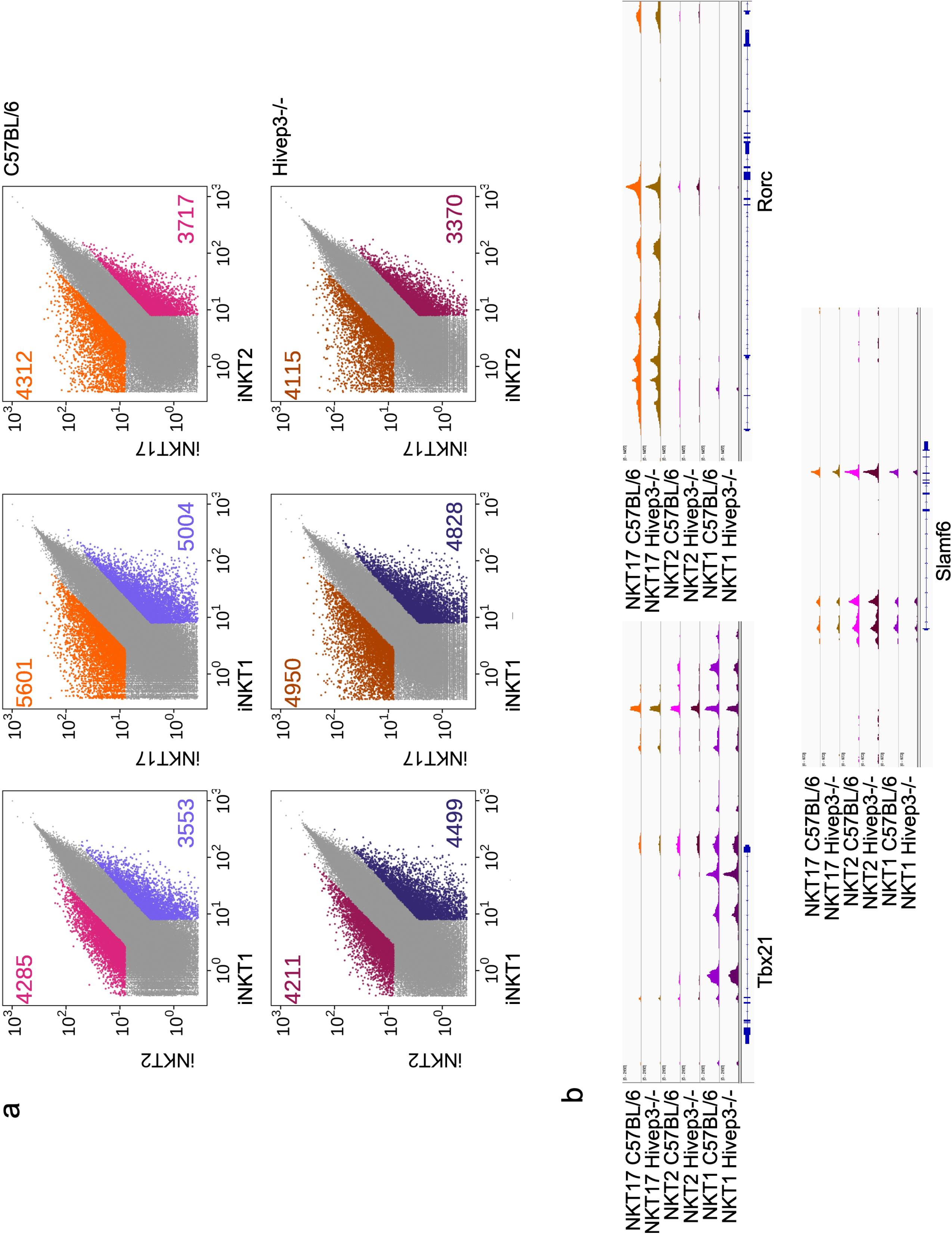
Genome-wide chromatin landscapes readily identify iNKT subsets in both C57BL/6 and *Hivep3*^−/−^ mice. (**a**) Scatterplots of mean ATAC-seq counts per peak comparing iNKT1, iNKT2 and iNKT17 subsets from C57BL/6 (top) or *Hivep3*^−/−^ (bottom) mice. Pairwise contrasts were performed with limma and differentially accessible regions (colored) were filtered based on an FDR adjusted p-value of less than 0.1 and an estimated fold-change of at least 3. (**b**) Mean ATAC-seq coverage at the *Tbx21*, *Rorc* and *Slamf6* loci for ATAC-Seq tracks.

**Supplementary Figure 7.**
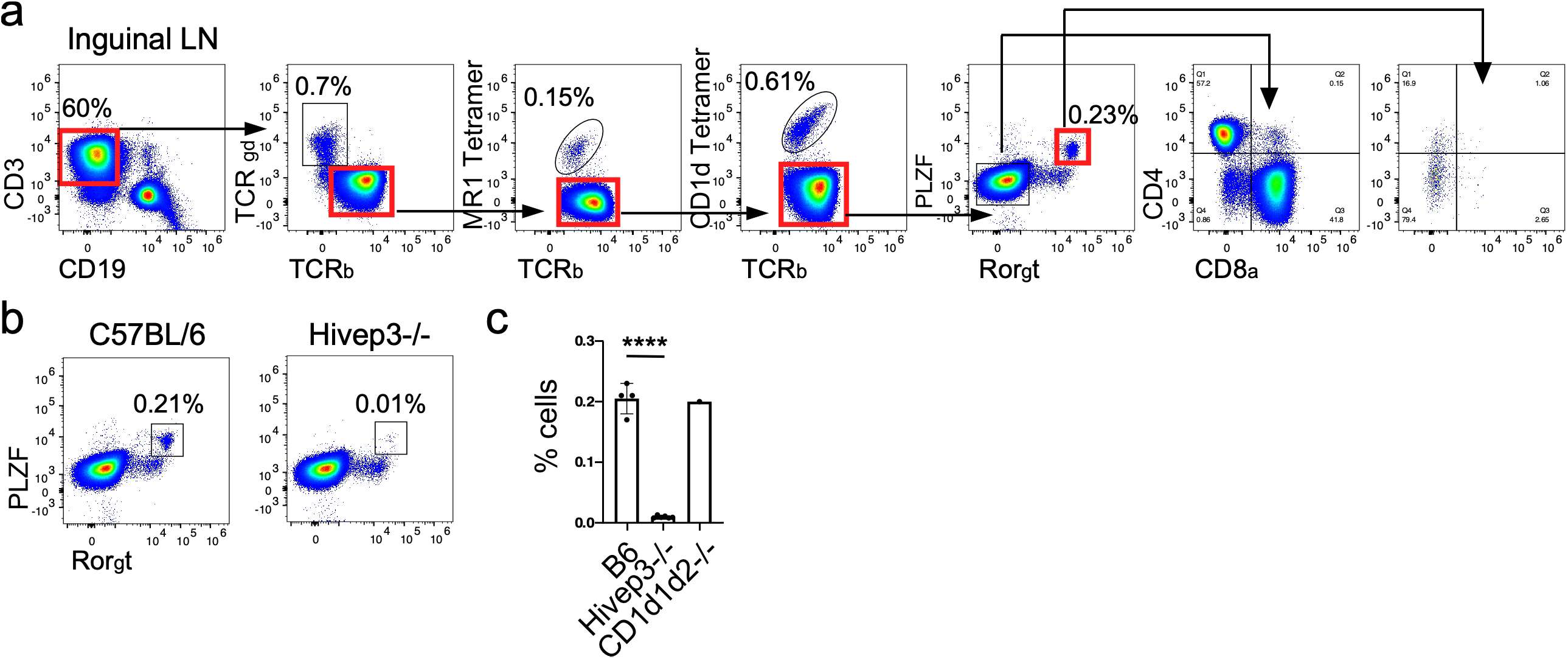
Identification of CD4^−^CD8^−^ TCRβ^+^ T cell population that co-express PLZF and Rorγt in the inguinal lymph nodes of mice. (**a**) Gating strategy used to identify non-iNKT, non-MAIT, αβ T cells with co-expression of PLZF and Rorγt. (**b**) Representative analysis of PLZF and Rorγt expression in non-iNKT, non-MAIT αβ T cells in the inguinal lymph nodes of C57BL/6 and Hivep3^−/−^ mice. (**c**) Quantified proportions of PLZF^+^Rorγt^+^ non-iNKT, non-MAIT αβ T cells in the inguinal lymph nodes of C57BL/6, *Hivep3*^−/−^ and CD1d1d2^−/−^ mice.

**Supplementary Figure 8.**
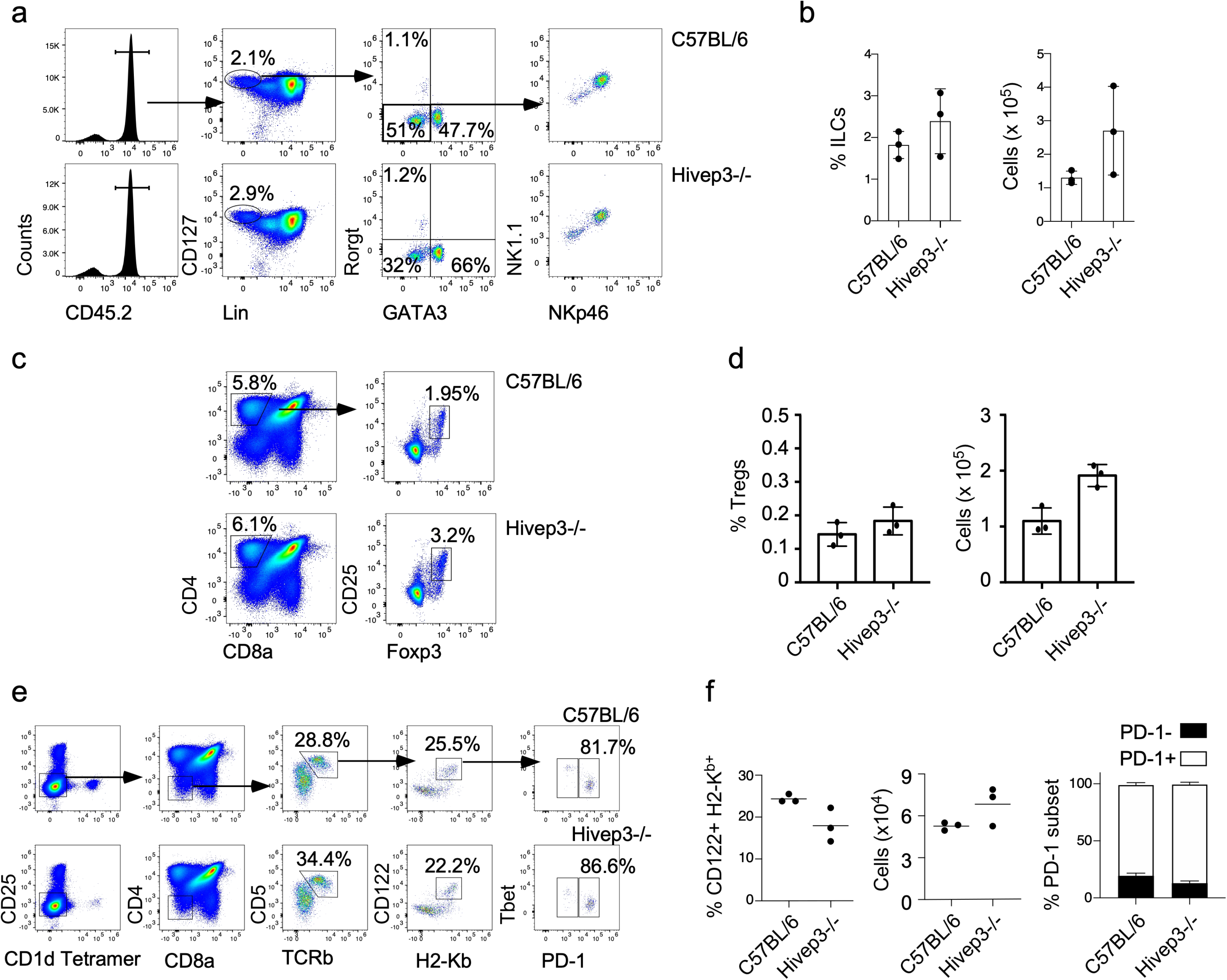
Analysis of innate lymphoid cells (ILCs), regulatory T cells and intraepithelial lymphocyte precursors in C57BL/6 and *Hivep3*^−/−^ mice. (**a**) Gating strategy used to identify and quantify the proportions of ILC populations in the lungs of C57BL/6 and *Hivep3*^−/−^ mice. (**b**) Quantified proportions and total numbers of ILCs in the lungs of C57BL/6 and *Hivep3*^−/−^ mice. (**c**) Gating strategy used to identify and quantify the proportions Tregs in the thymus of C57BL/6 and *Hivep3*^−/−^ mice. (**d**) Quantified proportions and total numbers of Tregs in the thymus of C57BL/6 and *Hivep3*^−/−^ mice. (**e**) Flow cytometry for the identification of IELp cells in the thymus. Numbers in plots indicate percent cells in outlined area. (**f**) summary of the frequency and number of mature CD122^+^ H-2K^b+^ cells gated as in (**e**) and total PD-1^+^ cells and PD-1^−^ in the thymic IELp population of C57BL/6 and *Hivep3*^−/−^ mice.

## Notes

### Competing Interest Statement

The authors have declared no competing interest.

